# Mechanism of gating and partial agonist action in the glycine receptor

**DOI:** 10.1101/786632

**Authors:** Jie Yu, Hongtao Zhu, Remigijus Lape, Timo Greiner, Rezvan Shahoei, Yuhang Wang, Juan Du, Wei Lü, Emad Tajkhorshid, Lucia Sivilotti, Eric Gouaux

**Author notes:** Van Andel Institute, 333 Bostwick Ave. NE, Grand Rapids, MI 49503, USA. These authors contributed equally (Jie Yu, Hongtao Zhu). Correspondence to Lucia Sivilotti or Eric Gouaux.

## Abstract

The glycine receptor is a pentameric, neurotransmitter-activated ion channel that transitions between closed/resting, open and desensitized states. Glycine, a full agonist, produces an open channel probability (Po) of ∼1.0 while partial agonists, such as taurine and γ-amino butyric acid (GABA) yield submaximal Po values. Despite extensive studies of pentameric Cys-loop receptors, there is little knowledge of the molecular mechanisms underpinning partial agonist action and how the receptor transitions from the closed to open and to desensitized conformations. Here we use electrophysiology and molecular dynamics (MD) simulations, together with a large ensemble of single particle cryo-EM reconstructions, to show how agonists populate agonist-bound yet closed channel states, thus explaining their lesser efficacy, yet also populate agonist-bound open and desensitized states. Measurements within the neurotransmitter binding pocket, as a function of bound agonist, provide a metric to correlate the extent of agonist-induced conformational changes to open channel probability across the Cys-loop receptor family.

## Introduction

Drugs that activate receptors (agonists) can differ in their effectiveness and produce different maximum responses on the same receptors^1^. This property is called efficacy by pharmacologists, and agonists with low efficacy are known as “partial agonists”, a term coined by Stephenson in 1956^2^. Early work by del Castillo and Katz^3^ on the muscle nicotinic acetylcholine ACh receptor (nAChR), a member of the pentameric ligand-gated channel superfamily (pLGIC), led to the hypothesis that efficacy is determined by the agonist’s ability to keep the channel open after it has bound. In LGICs, agonist efficacy is reflected in the maximum open probability seen in single-channel recordings when the channel is fully occupied by the agonist. Neurotransmitters are often full agonists^4, 5^, as exemplified by ACh on the muscle nAChR and glycine on the glycine receptor (GlyR), but partial agonists are of interest in therapeutics and include varenicline^6^, a partial agonist of neuronal nAChR, used as an aid to smoking cessation. Moreover, in the auditory system, GABA, a partial agonist of GlyRs, is co-released with glycine and both agonists act on the GlyR to sculpt receptor decay kinetics and the time course of postsynaptic responses^7^.

pLGICs include important therapeutic targets, such as neuronal and muscle nAChR, GABA receptors and 5-HT_3_ receptors (5-HT_3_R)^8^. Among pLGICs, GlyRs are an vehicle through which to investigate the mechanism of agonist efficacy because they can be studied by both single-molecule electrophysiological measurements of efficacy and high resolution structural studies. Recent electrophysiological studies of GlyRs showed that channel activation is not a simple isomerization from resting to open states, but rather involves a more complex landscape of one or more pre-open intermediates^4, 9,10^. The presence of multiple agonist-bound closed states is a general feature of the pLGIC superfamily and there is substantial evidence that the closed intermediate states stabilized by agonist binding have increased affinity for the agonist^11–14^. Fitting models that include activation intermediates to GlyR, nAChR and 5-HT_3_R single channel data have led to the proposal that the limited efficacy of partial agonists is due to their reduced ability to change the conformation to a short-lived pre-open intermediate (flipped/primed), rather than the reduced ability to open the receptor once the intermediate is reached^12, 15, 16^.

Here we investigate the structural basis of partial agonist activation in GlyR, a paradigm Cys-loop receptor, exploiting extensive functional characterization showing that taurine and GABA act as partial agonists^17–19^. Previous GlyR structures have shed light on the mechanism of action of full agonists and antagonists^20–23^, but structures of the receptor bound to partial agonists are still absent. Furthermore, the previously solved GlyR structures are in detergent micelles and lack the M3/M4 cytoplasmic loop, factors that likely underlie the anomalously high Po of the receptor constructs and the physiologically ‘too large’ open state of the ion channel pore ^24, 25^. Here we isolated the full length GlyR with native lipids and determined high resolution structures of the receptor bound to glycine, taurine or GABA. The receptors were imaged by cryo-electron microscopy (cryo-EM) in MSP nanodiscs or after solubilization by styrene maleic acid (SMA) copolymers^26^, showing by MD simulations that the SMA complex adopts a physiologically relevant open state. Importantly, cryo-EM structures of GlyR bound to taurine or GABA reveal, for the first time, agonist-bound closed states as well as open and desensitized states.

## Glycine-bound states

The wild-type GlyR is robustly activated by glycine and single-channel analysis of clusters of receptor activity, excluding the long-lived desensitized states, reveal a maximum open probability (P_O_) of 97% at saturating concentrations of glycine (Fig. 1a). Macroscopic current recordings of GlyR activity elicited by rapid application of 10 mM glycine to outside out patches show that the current decays to ∼ 57% of peak after 1s application because of desensitization (Extended Data Fig. 1a). Taken together, single channel and macroscopic results suggest that, under steady state conditions, the glycine-bound GlyR substantially populates both open and desensitized states. Hence, we reconstituted detergent-purified GlyR into nanodiscs or we directly extracted the receptor using SMA copolymers, followed by preparation of cryo-EM grids in the presence of 10 mM glycine.

**Figure 1.**
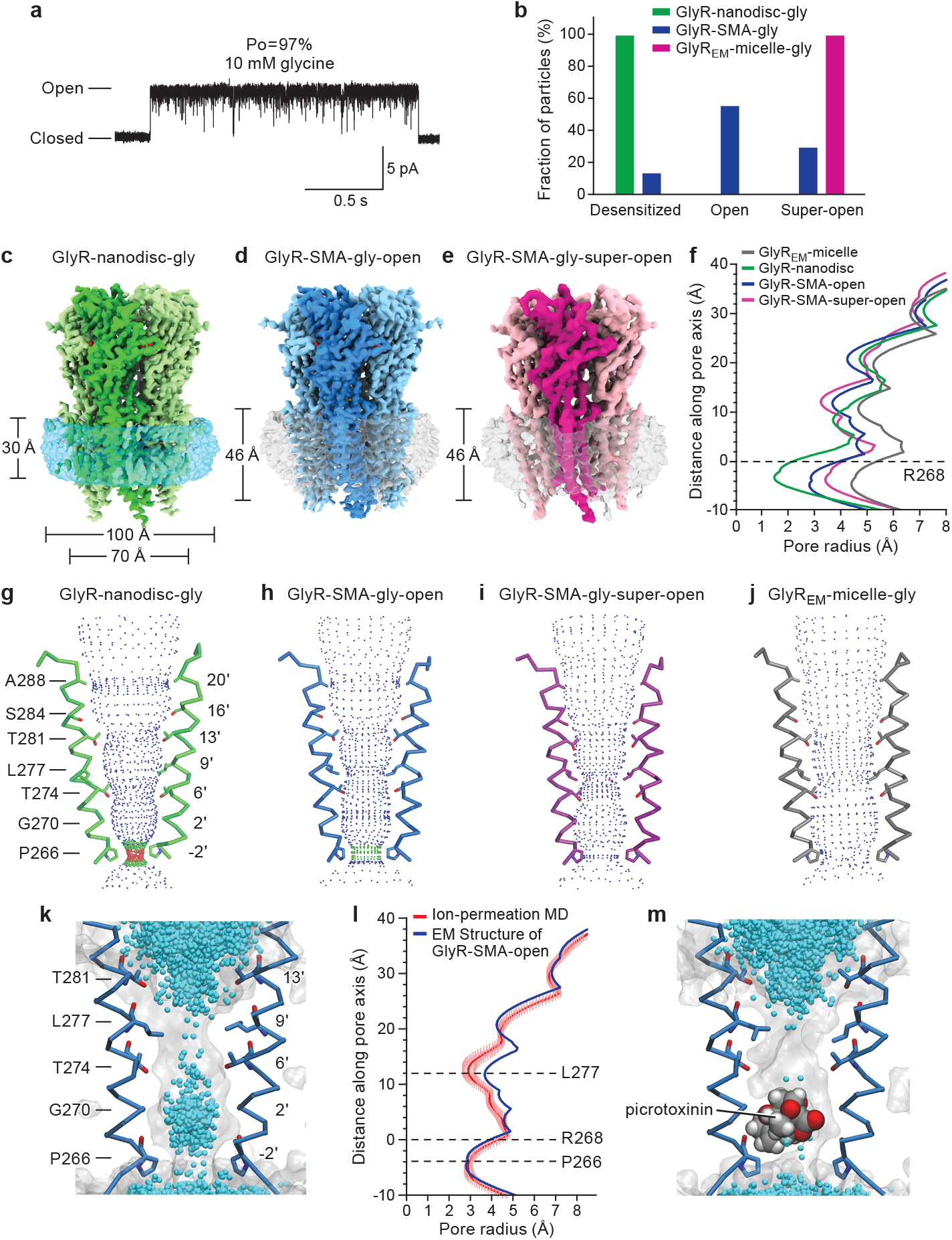
The GlyR bound to the full agonist glycine: different states and their pore profiles. **a,** Repres entative cell-attached single-channel recording of GlyR openings in the presence of 10 mM glycine. **b,** Bar chart showing the fraction of GlyR particles in the desensitized, open and super-open states in the nanodisc, SMA and micelle environments. **c,** Cryo-EM reconstruction map of GlyR complexed with glycine in the nanodisc environment, viewed parallel to the membrane (GlyR-nanodisc-gly). One subunit is highlighted in green. The nanodisc density is indicated with a blue transparent surface. **d-e,** Cryo-EM density maps of the open (GlyR-SMA-gly-open) and super-open (GlyR-SMA-gly-super-open) states of GlyR complexed with glycine in the SMA environment, with one subunit highlighted with blue and magenta, respectively. The transparent surface represents lipid-SMA density. **f,** Plots of pore radius as a function of distance along the pore axis for glycine-bound states in the micelle (GlyR_EM_-micelle, PDB code: 3JAE), nanodisc (GlyR-nanodisc) and SMA (GlyR-SMA-open and GlyR-SMA-super-open). The Cα position of Arg 268 (M2 0’) is set to 0. **g-j,** Ion permeation pathways for GlyR-nanodisc-gly (green), GlyR-SMA-gly-open (blue), GlyR-SMA-gly-super-open (magenta) and GlyR_EM_-micelle-gly (grey, PDB code: 3JAE). M2 helices of two subunits are shown in ribbon and the side chains of pore-lining residues are shown in sticks. As calculated by the program Hole, different colors define different radii: red <1.8 Å, green 1.8–3.3 Å, and blue > 3.3 Å. **k-m,** Snapshot from the ion-permeation molecular dynamics (MD) simulation of the GlyR-SMA-gly-open model at −100 mV without (k) or with the channel blocker picrotoxinin (m) placed in the pore. Permeating Cl^-^ ions from many frames of the simulation are shown as cyan spheres and the 50% water-occupancy isosurface is shown in transparent grey. The molecule of picrotoxinin is shown in van der Waals representation. **l,** Pore-radius profiles of structural models of the GlyR-SMA-gly-open. The profile of the cryo-EM structure model (in blue) is compared with the computed profile during the MD simulation (in red) illustrated in (k). The latter is displayed as the mean profile (darker line) ± 1 SD (lighter shade)—calculated from the different frames of the simulation.

The glycine-bound receptor in nanodiscs (GlyR-nanodisc-gly) yielded a reconstruction with a single 3D class at 3.2 Å resolution (Fig. 1c, Extended Data Fig. 7, Supplementary Fig. 1, Supplementary Table 1). The receptor features are well resolved and there is clear density for lipids and the membrane scaffold protein (MSP) (Fig. 1c). Within the receptor transmembrane domain (TMD), at the 9’L and −2’P constriction sites on the M2 helix, the pore has a radius of 4 Å and 1.5 Å, respectively, suggesting that the 9’L gate is ‘open’ and the −2’P gate is closed, consistent with the receptor occupying a desensitized-like conformation (Fig. 1f). The presence of the glycine-bound receptor in a single 3D class that is structurally a closed desensitized state is at odds with electrophysiology data. We thus reasoned that either the lipids or perhaps the nanodisc complex itself shifts the conformational equilibrium of the receptor to the desensitized state.

We proceeded to directly excise the receptor from cell membranes using SMA copolymers, isolating the receptor in its complex with endogenous lipids. Subsequent single particle cryo-EM studies revealed open (GlyR-SMA-gly-open), desensitized (GlyR-SMA-gly-desensitized) and super-open (GlyR-SMA-gly-super-open) states with overall resolutions of 2.9 Å, 3.1 Å and 4.0 Å, respectively (Figs. 1d-e, Extended Data Fig. 7, Supplementary Tables 2 and 5). Importantly, these reconstructions have well resolved transmembrane domain (TMD) densities, especially for the M2 helices, allowing us to precisely locate the 9’L and −2’P residues and, from these positions, define the functional state of each structure (Supplementary Fig. 4). In all of the structures the ion channel pores are open at 9’L but adopt three distinct conformations at the −2’P position (Fig. 1g-i, Extended Data Figs. 1b-c and 3).

The three distinct conformations at the −2’P position, in turn, represent open, desensitized and a super-open state, the latter of which is likely not physiologically relevant. In the GlyR-SMA-gly-open state the constriction of the pore is ∼5.6 Å, in agreement with the estimated pore diameter value of ∼5.34 Å and sufficient to allow permeation of partially hydrated chloride ions^27^ and block by cyanotriphenylborate^28^ (Fig. 1d, f). The GlyR-SMA-gly-desensitized state is similar to the GlyR-nanodisc-gly conformation, with a constriction at −2’P of 3 Å in diameter, indicative of a non-conducting state (Extended Data Fig. 1b, c). In the GlyR-SMA-gly-super-open state the pore is ∼7 Å in diameter at −2’P, a dimension that is larger than that estimated by electrophysiological sizing experiments^29^ but smaller than that of the truncated receptor in detergent micelles (GlyR_EM_-micelle-gly; PDB code: 3JAE)^20^ (Figs. 1f, i, j). Interestingly, we find density protrusions in the super-open conformation at −2’P yet, due to the moderate resolution of the reconstruction, are unable to accurately model the density (Extended Data Fig. 4). Most importantly, the SMA-solubilized receptor yields conformations in physiologically relevant open and desensitized states, approximately mirroring the population of states estimated from electrophysiological studies (Fig. 1b).

To rigorously assess the conductance of the three conformations of the glycine-bound receptor in SMA, we performed MD simulations. The GlyR-SMA-gly-open conformation is Cl^-^ permeable, with a calculated conductance of ∼45 pS, a value qualitatively comparable to the reported experimental values of 80-88 pS^30^ (Fig. 1k, l) and, unlike the GlyR_EM_-micelle-gly state, picrotoxinin blocks ion conductance^25^ (Fig. 1m). By contrast, no Cl^-^ permeation events were observed during the 500 ns simulation for the GlyR-SMA-gly-desensitized state, consistent with a non-conductive conformation. The GlyR-SMA-gly-super-open structure is unstable during MD simulations and collapses to a non-conductive channel (Extended Data Fig. 2b). We speculate that factors, such as specific lipids not present in the simulation, stabilize the super open state under experimental conditions and that the super open state is not physiologically relevant.

## Partial agonist bound states

Taurine, ubiquitous throughout the nervous system, and GABA, a crucial inhibitory neurotransmitter, act as partial agonists on the GlyR. Single-channel recordings of GlyR in the presence of 100 mM taurine or GABA, in comparison to glycine, feature additional long-lived shut states while the channel is not desensitized, especially for GABA, suggesting the presence of a substantial population of partial agonist-bound shut states (Fig. 2a). Indeed, the maximum P_O_ for taurine and GABA is 66% and 39%, respectively, substantially lower than that of glycine (97%; Fig. 1a). The partial agonists are also less potent, with effective concentrations (EC_50_) for taurine and GABA of 1050 ± 80 µM and 28.4 ± 0.9 mM, respectively (cf. glycine: 190 ± 20 µM; Fig. 2c). In the subsequent structural studies, we applied 20 mM taurine and 40 mM GABA to the receptor isolated in SMA or reconstituted into nanodiscs.

**Figure 2.**
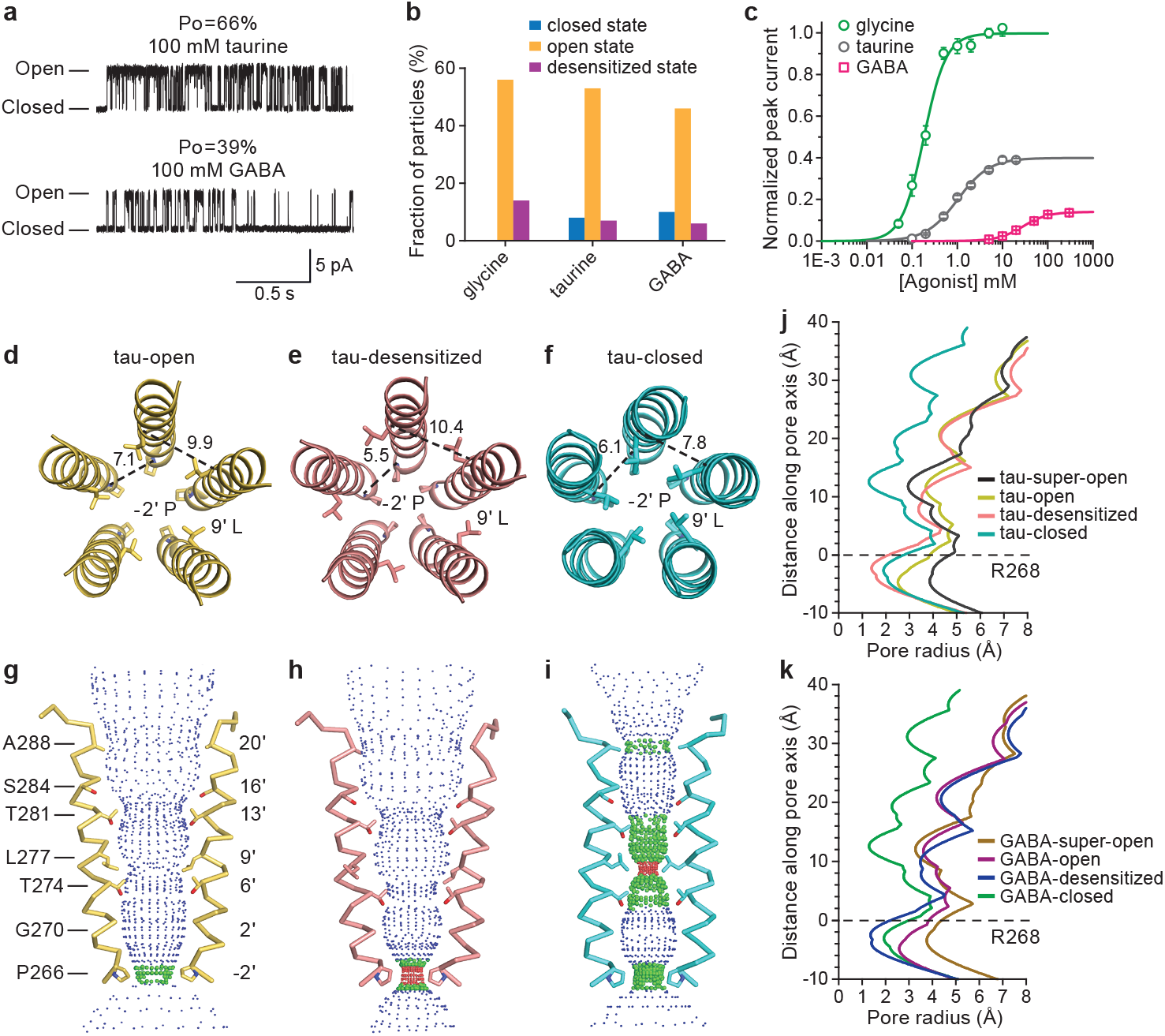
Partial agonist-bound states and ion channels. **a,** Representative cell-attached single-channel recordings of GlyR openings elicited by 100 mM taurine and 100 mM GABA, respectively. **b,** Fractions of GlyR particles in the open, closed and desensitized states for GlyR complexed with glycine, taurine and GABA and characterized in the SMA environment. Fraction of particles in the super-open state were not counted for clarity. **c,** Dose-response data for glycine, taurine and GABA from HEK 293 cells expressing GlyR. EC_50_ for glycine, taurine and GABA are 190 ± 20 µM, 1050 ± 80 µM and 28.4 ± 0.9 mM, respectively. Error bars represent SEM and n≥6 cells for all the experiments. Curves are normalized to the glycine maximum current recorded in each cell. **d-f,** Conformation of the residues −2’P and 9’L at the main constriction sites in the taurine-bound GlyR at the open (tau-open, yellow), desensitized (tau-desensitized, salmon) and closed (tau-closed, cyan) states in the SMA. The five M2 helices in each state are shown as cartoon and the side chains of −2’P and 9’ L are shown in sticks. The Cα distances between two adjacent −2’P or 9’L are denoted in angstrom. **g-i,** Ion permeation pathway for taurine-bound states in the SMA. M2 helices of two subunits are shown in ribbon and the side chains of pore-lining residues are shown in sticks. As calculated by the program Hole, different colors define different radii: red <1.8 Å, green 1.8–3.3 Å, and blue > 3.3 Å. **j,** Plots for pore radius as a function of distance along the pore axis for taurine-bound states in the SMA. The Cα position of Arg 268 is set to 0. **k,** Plots for pore radius as a function of distance along the pore axis for GABA-bound GlyR at the closed (GABA-closed), open (GABA-open), desensitized (GABA-desensitized) and super-open (GABA-super-open) states in the SMA. The Cα position of Arg 268 is set to 0.

Three-dimensional classification of the SMA and nanodisc cryo-EM data sets, for each partial agonist, reveals 4 and 2 distinct classes, respectively, with overall resolutions of 2.8 to 3.3 Å and local resolutions as high as 2.5 Å in the extracellular domain (ECD) (Extended Data Fig. 8 and 9, Supplementary Tables 1-3 and 5, Supplementary Fig. 2, 4, 5). In the SMA data sets, three of the four taurine-or GABA-bound structures are overall similar to the glycine-bound structures and correspond to open, desensitized, and super-open states (Fig. 2d-k). By contrast, in the nanodisc data sets, one of the two structures is reminiscent of the glycine-bound desensitized state and we did not observe the open and super-open states as in the SMA data sets, supporting the notion that native lipids extracted by SMA best enable the receptor to occupy physiologically relevant states (Extended Data Fig. 1d, e).

We discovered a previously unseen, partial agonist-bound conformation of the receptor, observed with taurine and GABA, and in both the SMA preparation and the receptor in nanodiscs. In these reconstructions, there is clear density for the partial agonists in the orthosteric binding pocket (Supplementary figure 6) and the M2 helices are oriented approximately perpendicular to the membrane. In these partial agonist-bound structures, the pore is closed, as the side chains of 9’L point to the center of the channel, creating a constriction less than 3 Å in diameter and thus rendering the channel impermeable to Cl^-^ (Fig. 2f, i, j, k, Extended Data Fig. 1d-f). At the −2’P site, the pore has a diameter of 4 Å, intermediate between the open and desensitized states (Fig. 2j-k, Extended Data Fig. 1f).

The closed state observed in the partial agonist-bound cryo-EM datasets provides the first structural visualization of the long-lived shut states in single-channel recordings, and is consistent with these shut states being most highly populated in the presence of partial agonists (Fig. 2a). By contrast, the glycine-bound closed conformation of the receptor is short-lived, as the receptor rapidly and nearly completely transitions to the brief intermediate “flipped” state^12^, thus precluding its capture by single-particle cryo-EM. Taurine or GABA’s less potent ability to ‘flip’ GlyR, as well as their propensity to promote the receptor’s return from the flipped state to the closed state, allows us to capture agonist-bound closed states via cryo-EM. We suggest that the cryo-EM partial agonist closed states, bound with either GABA or taurine, most likely represent agonist-bound, unflipped states. In each cryo-EM dataset, the fraction of particles in each state is roughly related to the likelihood of the receptor occupying that conformation. The particle fractions of the open or desensitized states decrease for taurine as compared to glycine, and decrease further for GABA. Conversely, the fraction of particles in the closed state is highest for the GABA dataset, lower for the taurine dataset, and absent in the glycine dataset (Fig. 2b). These findings are consistent with single-channel recordings, showing GABA having the lowest efficacy for activating GlyR, followed by taurine, with glycine having the highest efficacy.

## Neurotransmitter binding site

In all complexes, the densities for glycine, taurine and GABA are distinctive and well defined, enabling accurate definition of the positions and poses of the agonists (Fig. 3a-c, Extended Data Figs. 7, 8 and 9). In the glycine complex, the agonist’s amino group forms hydrogen bonds with the backbone carbonyl oxygen of Phe175 (loop B) and a cation-π interaction with Phe223 (loop C), while the carboxyl group interacts with Thr220 (loop C), Arg81 (β2) and Ser145 (β6) (Fig. 3d), consistent with the GlyR-α3 crystal structure^22^. While the orientation of taurine and GABA is similar to that of glycine, the interactions between the amino groups of the agonists and receptor residues are different (Fig. 3e-f). In the taurine complex, an additional hydrogen bond is formed between the amino group and the backbone carbonyl oxygen of Ser174 whereas no cation-π interactions with Phe223 were identified (Fig. 3e). In the GABA complex, the amino group forms hydrogen bonds with the backbone carbonyl oxygen of Ser174 and Glu173 (loop B) (Fig. 3f). As in the glycine complex, the sulfate group of taurine and the carboxylate group of GABA participate in hydrogen bonds with Thr220, Arg81 and Ser145.

**Figure 3.**
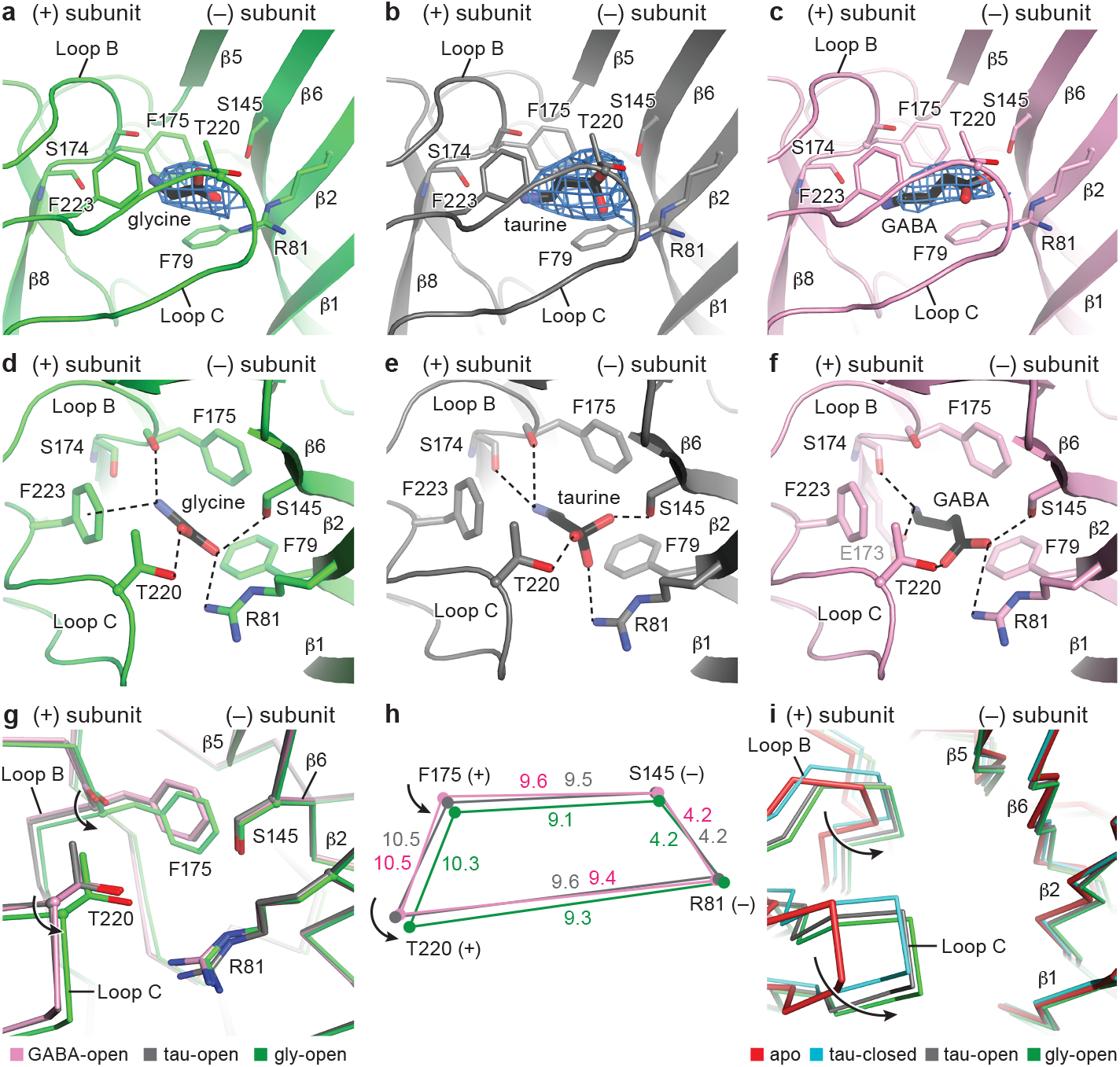
Neurotransmitter binding sites. **a-c**, Illustration of the densities contributed by the glycine, taurine and GABA, contoured at 11 σ, 10 σ, and 8 σ, respectively. For the neurotransmitters, carbon and sulfate atoms are colored in black, whereas nitrogen and oxygen atoms are in blue and red, respectively. **d-f,** Glycine, taurine and GABA binding sites with possible hydrogen bonds and cation-π interactions shown as dashed lines. **g,** Comparison of the conformational changes produced by the binding of glycine (green), taurine (grey) and GABA (pink) to the agonist site detected by superimposing the extracellular domains (ECDs) among the glycine, taurine and GABA-bound open states. The four key residues are indicated and shown in sticks with the Cα atoms represented in spheres. **h,** Schematic diagram illustrating the changes in distances of the Cα atoms of R81, F175, S145 and T220 in the glycine, taurine and GABA-bound open states. The values colored in green, grey and pink represent the measured distances in angstrom with glycine, taurine and GABA in the binding site. **i,** Conformational changes of the binding pockets in the apo state (red), glycine-bound open (forest), taurine-bound open (grey) and taurine-bound closed (cyan) states by superimposing the main chain atoms of the ECDs.

To identify the changes in the binding pocket upon agonist binding, we solved the structure of a functional receptor variant with a truncated M3/M4 loop in the apo/resting state (apo-GlyR_EM_; Supplementary Figure 3, Supplementary Table 4). Because the M3/M4 loop is far from the agonist binding site, and because the apo receptor is in a closed/resting state conformation, the loop deletion does not substantially affect receptor conformation in the apo state. Indeed, the apo-GlyR_EM_ structure has a pore constriction at 9’Leu, consistent with the apo-GluCl^31^ and taurine/GABA-bound closed sate (Extended Data Fig. 1f). Upon comparison of the apo/resting state conformation of the agonist binding pocket with the open state/agonist-bound complexes, the binding pocket has a profound ‘contraction’ that largely involves loop C transitioning from an “uncapped” to “capped” configuration and loop B moving towards to the agonist (Fig. 3i), reminiscent of conformational changes in the acetylcholine receptors^14, 32, 33^.

The extent of binding pocket contraction, however, is different between full and partial agonists, with the volume of the agonist binding pocket smallest in the glycine-bound structure. Thr220 in loop C, Phe175 in loop B, Arg81 in β2, and Ser145 in β6 all coordinate taurine and GABA similarly, yet slightly different than their coordination of glycine. In the glycine complex, the Cα atoms of Thr220 in loop C and Phe175 in loop B ‘contract’ toward the agonist by 0.5 Å and 0.6 Å, respectively, compared with the taurine-bound pocket (Fig. 3g, h). We propose that these conformational changes are coupled to agonist-induced activation^29^, consistent with the notion that the agonist has an increased stabilization of the activated configuration^34^. Thus, in the full agonist open state, loops B and C move closer to the ligand in comparison to partial agonists, resulting in a smaller binding pocket and a more efficient activation of the receptor (Fig. 3h). Our structural data, together with agonist-dependent measurements on the AChR, 5HT-3 and GABA receptor, show that full and partial agonist complexes of the open state of a Cys-loop receptor, within the confines of the agonist-binding site, are not identical and that full agonists yield a more ‘compact’ binding site in comparison to partial agonists (Supplementary Table 6).

## GABA acts on YGF mutant with a high Po

To test if contraction of the agonist binding pocket by partial agonists is correlated to agonist efficacy, we investigated the F175Y/Y177F variant (YGF) ^19^, a double mutant that involves only a single oxygen atom swap from one residue to another. Remarkably, GABA is an almost full agonist on the YGF mutant, with a Po of ∼96% (Fig. 4b). In the mutant, the potency of GABA in activating the channel (EC_50_) and in displacing [H^3^]-strychnine (Ki) also increases by 15- and 8-fold, respectively. Importantly, glycine remains a full agonist, with a Po of nearly 1.0, and the 5-fold decrease in its EC_50_ suggests glycine efficacy is also increased (Fig. 4a). To explore the structural basis for the increase in GABA potency, we solved the cryo-EM structure of the YGF mutant bound with GABA in SMA, obtaining high-resolution maps corresponding to the open, desensitized and super-open states (Fig. 4h, Extended Data Fig. 10, Supplementary Fig. 5, Supplementary Table 4). Importantly, we did not observe any 3D classes associated with the closed state of the receptor, corroborating the electrophysiological data showing that GABA acts as a nearly full agonist on the YGF mutant.

**Figure 4.**
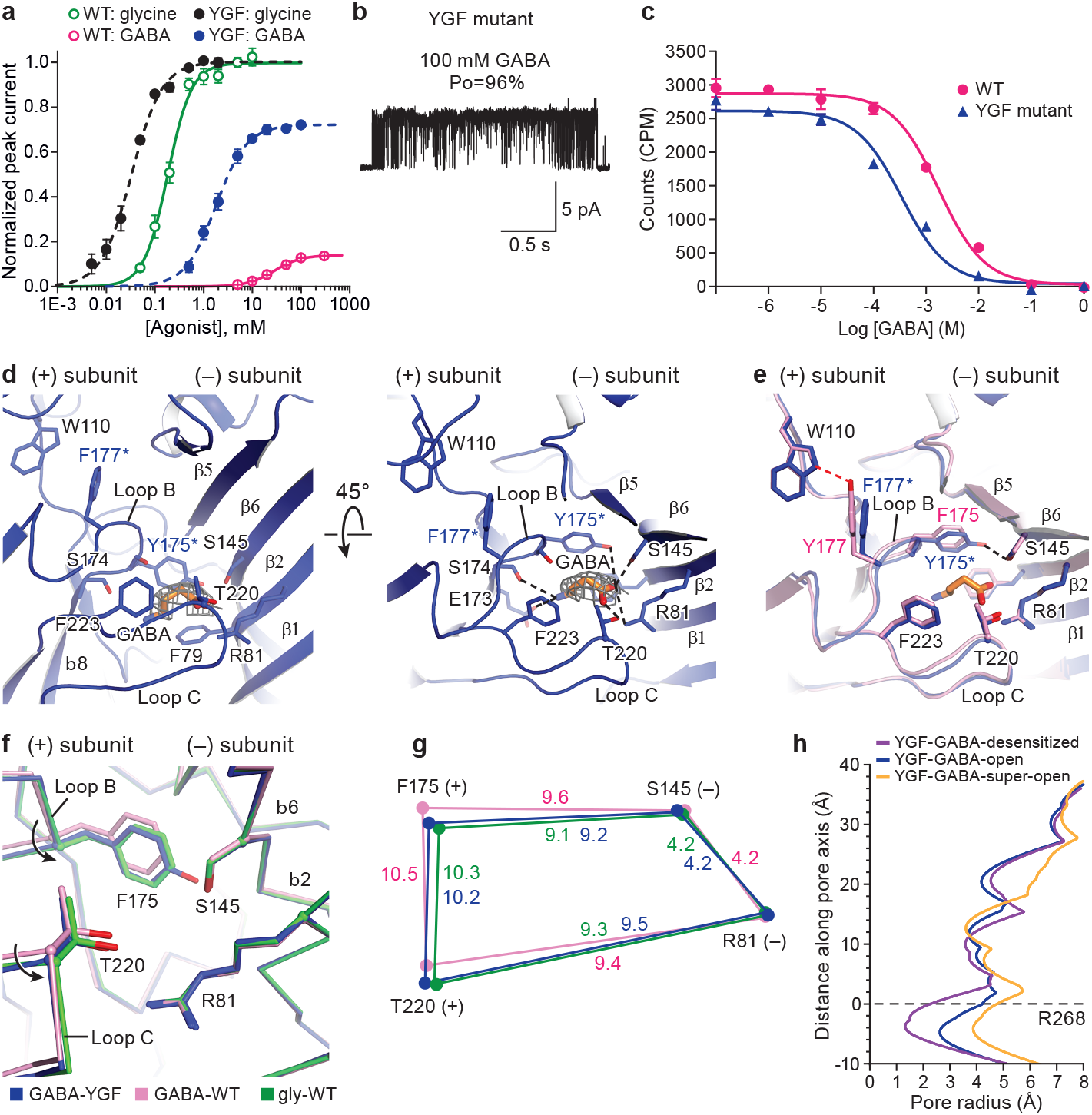
The YGF mutation in loop B makes GABA a high efficacy agonist. **a,** Dose-response data for glycine and GABA from HEK293 cells expressing wild type (WT) GlyR receptor or YGF mutant. EC_50_ for glycine to WT and YGF mutant are 190 ± 20 µM and 33 ± 3 µM, respectively. EC_50_ for GABA to WT and YGF mutant are 28.4 ± 0.9 mM and 1.05 ± 0.08 mM, respectively. Error bars represent SEM and n≥6 cells for all the experiments. Responses are normalized to maximum responses to glycine in each cell. Glycine and GABA have a higher efficacy in the YGF mutant. **b,** Cell-attached single-channel recording showing openings of the YGF mutant produced by 100 mM GABA.. **c,** Competition ligand binding experiment where the binding of ^3^H-strychnine is displaced by cold GABA in the WT and YGF mutant. Data are shown as means ± SEM (n=3). Ki of GABA to WT and YGF mutant are 1.41 ± 0.94 mM and 0.18 ± 0.07 mM, respectively. **d,** GABA binding site in the YGF mutant viewed parallel to the membrane (left) or from the extracellular side of the membrane (right). GABA density is contoured at 0.013 σ. The possible hydrogen bonds and cation-π interactions are shown as dashed lines. **e,** Superimposition of the binding pockets from the WT and YGF mutant in the presence of GABA to show the impact on the binding pocket of the swap between Y177 and F175. **f,** Conformational changes in the binding pockets of the open states of GABA-bound YGF mutant (blue) and WT (pink) and glycine-bound WT (green) shown by superposing the ECDs. The four key residues are indicated and shown in sticks with the Cα atoms represented in spheres. **g,** Schematic diagram illustrating changes in the distances of the Cα atoms of R81, F175, S145 and T220 in the open states of GABA-bound YGF mutant (blue) and WT (pink) as well as glycine-bound WT (green). **h,** Plots for pore radius as a function of distance along the pore axis for GABA-bound YGF mutant in the open, desensitized and super-open states. The Cα position of Arg 268 is set to 0.

The increased binding affinity of GABA to the YGF mutant arises from additional interactions involving the carboxylate group of GABA, including a hydrogen bond with the hydroxyl oxygen of Tyr175 (Fig. 4d). Superposition of the binding pockets of the WT and YGF mutant highlights other differences between interactions by receptor side chains. The mutation of Tyr177 to Phe disrupts the hydrogen bond formed with Trp110 through the hydroxyl oxygen of Tyr177, whereas the mutation of Phe175 to Tyr yields an additional hydrogen bond with Ser145 (Fig. 4e). These new receptor interactions enable loop B, on which Tyr175 and Phe177 reside, to move closer to the (-) subunit, reducing the binding pocket volume to a value similar to that of the WT glycine open complex (Fig. 4f, g). Thus, binding pocket volume is a key determinant of agonist efficacy, and contraction in the binding pocket facilitates transition of the receptor from the resting state to the flipped and, ultimately, the open state.

To understand why GABA is almost a full agonist at the GABA receptor (GABAR) but a partial agonist for GlyR, we also superimposed the binding pocket of GlyR with the canonical GABA binding site of the GABAR^35, 36^. Remarkably, most of the residues coordinating GABA in the two receptors are identical, with the exception of Tyr97, Tyr157, and Tyr205 in GABAR, which are Phe115, Phe175 and Phe223 in GlyR, respectively (Extended Data Fig. 5a, b). Moreover, the binding pocket of the GlyR open state has a smaller volume than GABA-bound GABAR (Extended Data Fig. 5c), implying that the opening of GABAR requires a smaller contraction of the binding pocket to fully transition the receptor to the open state. To explore if the simple substitution of the three tyrosine residues of GABAR with the phenylalanine residues of GlyR enables GABA to act as a full agonist on the GlyR, we prepared the YYY mutant (F115Y/F175Y/F223Y) and found that GABA remained a weak partial agonist, with a similar Po in the YYY mutant as in the WT receptor (Extended Data Fig. 5d). Thus, the tuning of agonist efficacy requires precise changes in the pocket volume and in agonist-receptor interactions that go beyond which residues interact directly with the agonist.

## ECD and ECD-TMD interface changes upon binding to partial agonists

The transition from the partial agonist-bound closed state to the open state involves substantial conformational changes. Because the taurine-bound structures are most well resolved, and similar to the GABA-bound structures, we used the taurine SMA complexes for analysis of the ECD and ECD-TMD interfaces. Upon the closed-to-open transition, Phe175 in loop B and Thr220 in loop C move towards the agonist by 0.7 Å and 0.8 Å, respectively (Fig. 5a, c, f). Additionally, the Cα atoms of Ser145 in the β6 and Arg81 in the β2 shift away from the (+) subunit by 0.5 Å and 0.6 Å, respectively. Transduction of the conformational changes from the agonist binding pocket to the ECD-TMD interfaces include the repositioning of the β8-β9 loop by loop C and loop B, reshaping of β1-β2 loop by β2, reseating of the Cys-loop by β6 and loop B, and movement of pre-M1 by loop C (Fig. 5e and Extended Data Fig. 2a). The M2-M3 loop acts as a “bridge”, forming contacts with the ECD and repositioning the M2 and M3 helices. To quantify the movement in the M2-M3 loop ‘against’ the ECD, we measured the distance between Thr70 in the β1-β2 loop and Pro291 in the M2-M3 loop, and between Gln202 in the β8-β9 loop and Ser294 in the M2-M3 loop. From the tau-closed to the tau-open state, the M2-M3 loop moves away from the ion channel pore, resulting in a decrease of 2.4 Å in the distance between Thr70 and Pro291, and an increase of 2.4 Å in the distance between Gln202 and Ser294 (Fig. 5g). Pro291, positioned beneath Thr70, stabilizes the M2 and M3 helices in the tau-open state, reminiscent of the position of Val45 and Pro268 in the GluCl in complex with ivermectin^37^. Furthermore in the tau-closed state, a hydrogen bond between the side chain of Ser294 and the backbone amino group of Gln235 is ruptured during receptor activation, facilitating the outward movement of the M2 helix (Fig. 5e, g). This action agrees with reports that that mutation of Ser294 to cysteine increases glycine EC50 by ∼5 fold^38^.

**Figure 5.**
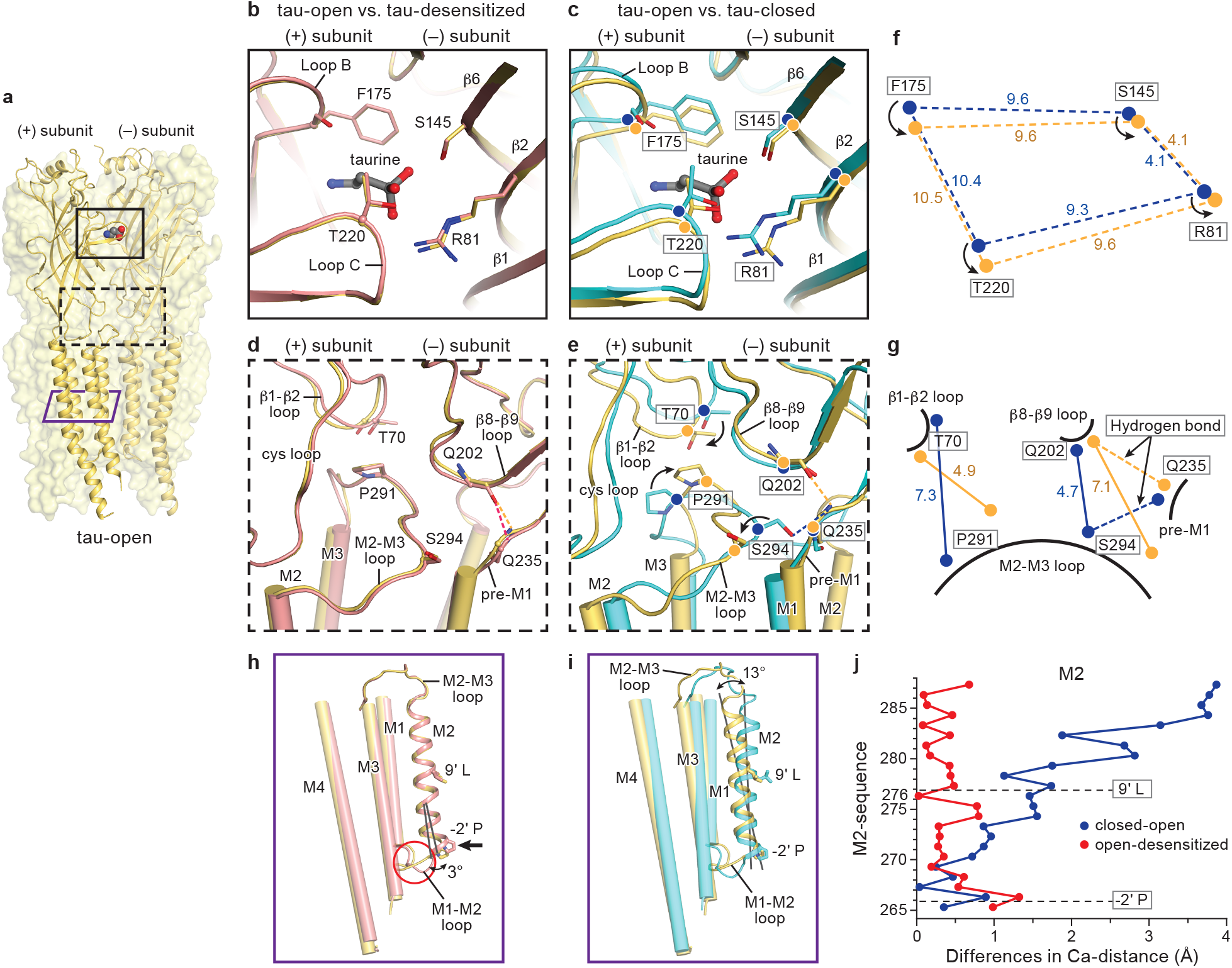
Conformational changes between the open (tau-open), desensitized (tau-desensitized) and closed (tau-closed) states of taurine-bound GlyR in the SMA. **a,** Reference orientation of tau-open state. Two of subunits are shown as cartoon and the other three subunits are represented as surface. The partial agonist taurine is highlighted by spheres. The binding pocket in the ECD, ECD-TMD interface and TMD are indicated by a black solid, a black dashed and a purple solid rectangle. **b-g,** Superposition of the (-) subunit to see the relative movements in the (+) subunit. The ligand taurine is shown in sticks. The key residues and secondary structures are noted. Hydrogen bonds are shown in dashed lines. The tau-open, tau-desensitized and tau-closed are colored in yellow, salmon and cyan, respectively. The Cα atoms of important residues in the tau-open and tau-closed state are highlighted by yellow and blue balls, respectively. (b,d) Comparison of the binding pocket (b) and ECD-TMD interfaces (d) between tau-open and tau-desensitized states. (c,e) Conformational changes in the binding pockets (c) and the ECD-TMD interfaces (f) between tau-closed, tau-open states. (f,g) Schematic diagrams illustrating the changes in distances of the Cα atoms of key residues in the binding pockets (f) and ECD-TMD interfaces (g) between tau-open and tau-closed states. Cα atoms in the tau-open and tau-closed states are represented by yellow and blue balls, respectively. **h-i,** Superimposition of a single subunit to see the changes in the TMD between tau-closed, tau-open and tau-desensitized states. M1, M3 and M4 helices are shown in cylinder while M2 helix is shown in cartoon. Two constriction sites (9’L and −2’P) are shown in sticks. The relative rotation angles of the M2 helix are indicated. When going from tau-open to tau-desensitized states, changes are concentrated on the M2 helix and M1-M2 loop (h). Larger movements, involving the whole TMD, occur in the transition from tau-closed to tau-open states (i). **j,** The plot of the differences in the position of Cα atoms from residues in the M2 helix derived from the tau-open, tau-desensitized and tau-closed states. Two constriction sites are indicated by dash lines.

In contrast with the transition from the closed to the open state, the tau-open to the tau-desensitized state shows that both the binding pockets and the ECD-TMD interfaces adopt the same conformation (Fig. 5a, b, d). This confirms previous hypotheses that in the open and desensitized states the binding pocket is in the same configuration and has the same affinity for the agonist. The transition from open to desensitized states may involve only local interactions in the ion permeation pathway^39, 40^, with desensitization proceeding from an uncoupling between the ECD and TMD^41^. In the ECD-TMD interfaces, a backbone hydrogen-bond between the Gln235 in the pre-M1 and Gln202 in the β8-β9 loop stabilizes the receptor in the open and desensitized states (Fig. 5d). Consistent with our results, functional experiments have shown that mutation of Gln202 to residues that cannot preserve this hydrogen bond network, as in the *shaky* startle disease mouse, reduces glycine potency^42^.

In comparison to previously solved Cys-loop receptors in closed/resting states ^20, 21, 43–45^, the tau-closed state exhibits unique features. We compared the ECD of the tau-closed state with apo-GlyR_EM_ and apo 5-HT_3A_R to investigate the conformational changes in the ECD and ECD-TMD interfaces (Extended Data Fig. 6a-e). In the binding pockets, loop C in the tau-closed state is in a “capped” configuration as compared to the other two structures, likely due to the binding of agonist, further supporting our view that the tau-closed is distinct from the resting state (Extended Data Fig. 6a-b). While the ligand binding pocket shows substantial conformational differences between the apo-GlyR_EM_ and tau-closed structures, the ECD-TMD interfaces of the two structures are remarkably similar, with identical conformations for M2-M3, β1-β2, β8-β9 and Cys loops (Extended Data Fig. 6d), suggesting that local movements of loop C alone cannot induce substantial conformational changes that affect the receptor ECD-TMD interfaces as well as the TMDs. Much like the ligand binding pocket, the 5-HT_3A_R ECD-TMD interface shows profound differences compared with the tau-closed state (Extended Data Fig. 6c, e).

## Conformational changes of the ion channel pore

To visualize the conformational changes at the TMD coupled with the movements of ECD and ECD-TMD interfaces, we compared the TMDs from tau-closed, tau-open and tau-desensitized states. In the transition from the closed to the open states, the whole TMD of each subunit undergoes a counterclockwise outward rotation of ∼8.6 ° relative to the pore axis, viewed from the extracellular side, expanding the pore size by moving the Cα atoms of 9’L and −2’P by 2 Å and 1 Å, respectively, opening the ion channel pore and allowing for the passage of Cl^-^ (Fig. 5a, i, j and Extended Data Fig. 6f). Among the M1-M4 helices, the M2 helices have a more pronounced movement, with a rotation of ∼13°. During the transition from the open to the desensitized state, the major conformational changes are in the TMD, with each subunit undergoing a clockwise rotation by ∼2° (Extended Data Fig. 6g). As a consequence, although the extracellular upper half of the pore has a small movement at 9’L, the intracellular lower half of the pore is occluded at −2’P, moving by ∼1.4 Å, in agreement with observations in the homomeric-GABA_A_ receptor and the α4β2 nicotinic receptor in the desensitized state ^46–48^ (Fig. 5a, h, j). In this process, the M2 helix exhibits a larger motion, with an inward rotation of ∼3°. Notably, the M1-M2 loop undergoes a relatively large displacement during desensitization, in agreement with studies showing that mutation of the M1-M2 loop can strongly affect entry into desensitization^49, 50^.

## Partial agonist gating mechanism

Previously unseen partial agonist-bound closed states, poised between the resting and flipped state, have now been captured, in accord with the observed long-lived shut state in the single-channel recordings (Fig. 2a). Binding of the partial agonist induces a contraction in the binding pocket, yet to a lesser extent than the full agonist (Fig. 6a-c). These ECD changes promote the receptor transition from the closed to open state, in which the TMD undergoes an outward rotation and expands the channel pore at the −2’P and 9’L positions (Fig. 6c). Sustained binding of agonist promotes a further transition into the desensitized state, involving an inward rotation of the TMD and occlusion of the pore at −2’P (Fig. 6d). The capturing of the agonist-bound closed, open and desensitized states provides a structural basis of the partial agonist gating mechanism for GlyR as well as other members of the Cys-loop family, which can guide the development of new therapeutics.

**Figure 6.**
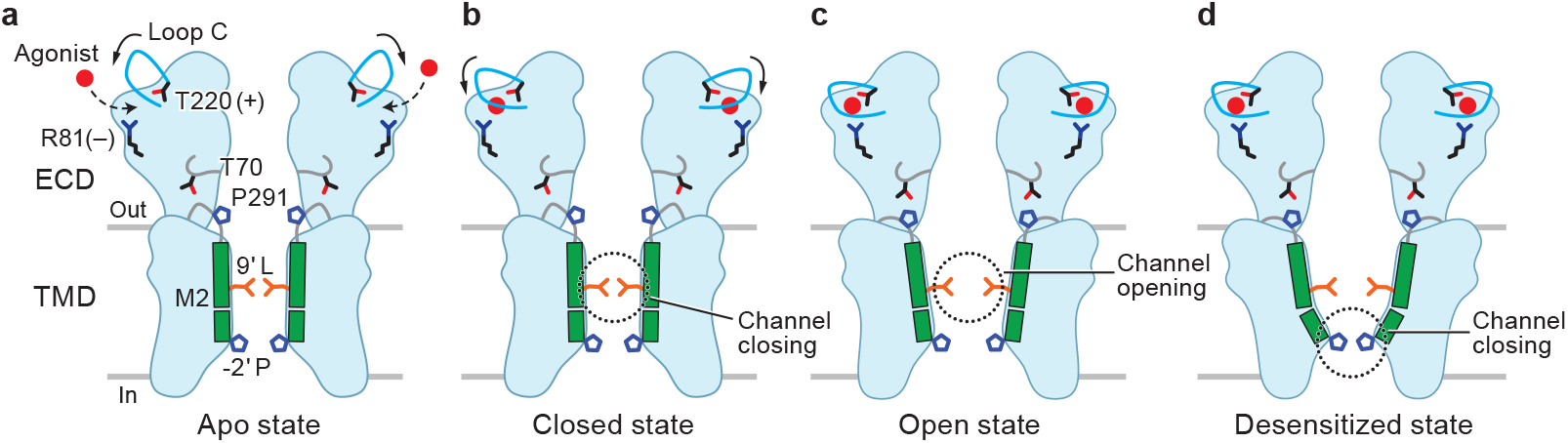
Cartoon models illustrating the gating mechanism of partial agonists. Important residues and two M2 helices are highlighted to show the conformational changes of the apo (a), closed (b), open (c) and desensitized (d) states, respectively. The agonist is shown as a red ball. The binding of partial agonist induces a contraction in the binding pocket, yet the channel is shut (a-b). The ECD changes promote the transition from closed to the open state (b-c), while sustained binding of partial agonist promote a further transition into the desensitized state (c-d).

## Acknowledgements

We thank L. Vaskalis for assistance with figures and F.Jalali-Yazdi for helpful discussions. We thank Polyscope for providing the XIRAN SL 30010 as a gift. Electron microscopy was performed at Oregon Health & Science University (OHSU) at the Multiscale Microscopy Core with technical support from the OHSU-FEI Living Lab and OHSU Center for Spatial Systems Biomedicine. Molecular simulations were performed using supercomputers from Blue Waters (ACI-1713784 to E.T.) and XSEDE (TG-MCA06N060 to E.T.). Blue Waters is supported by the National Science Foundation (OCI-0725070 and ACI-1238993) and the State of Illinois. XSEDE is supported by the National Science Foundation (ACI-1548562). This work was also supported by NIH grants P41-GM104601, U54-GM087519, and R01-GM123455 to E.T and R01-GM100400 to E.G. E.G. is an investigator of the Howard Hughes Medical Institute.

## Author contributions

J.Y., H.Z. and E.G. designed the project. J.Y. performed the sample preparation for cryo-EM and biochemistry studies. J.Y. and H.Z. performed the cyo-EM data collection. H.Z. and J.Y. analyzed the data. H.Z. performed the model building and ligand binding assay. J.D. and W.L. performed the cryo-EM data collection and processing for apo-GlyR_EM_. T.G., R.L. and L.G.S. performed the electrophysiological experiments and wrote the related method section. R.S., Y.W., and E.T. performed the MD simulations and wrote sections related to computational methods.

## Data Availability Statement

The data that support the findings of this study are available from the corresponding author upon request. The coordinates and associated volumes for the cryo-EM reconstruction of GlyR-gly-desensitized-nanodisc, GlyR-tau-closed-nanodisc, GlyR-tau-desensitized-nanodisc, GlyR-GABA-closed-nanodisc, GlyR-GABA-desensitized-nanodisc data sets have been deposited in the Protein Data Bank (PDB) and Electron Microscopy Data Bank (EMDB) under the accession codes 6PLR and 20373, 6PLT and 20375, 6PLS and 20374, 6PLV and 20377, and 6PLU and 20376. The coordinates and associated volumes for the cryo-EM reconstruction of GlyR-gly-open-SMA, GlyR-gly-desensitized-SMA and GlyR-gly-super-open-SMA data sets have been deposited in the PDB and EMDB under the accession codes 6PM6 and 20389, 6PM5 and 20388, and 6PM4 and 20386. The coordinates and associated volumes for the cryo-EM reconstruction of GlyR-tau-closed-SMA, GlyR-tau-open-SMA, GlyR-tau-desensitized-SMA and GlyR-tau-super-open-SMA data sets have been deposited in the PDB and EMDB under the accession codes 6PM3 and 20385, 6PM2 and 20384, 6PM1 and 20383 and 6PM0 and 20382. The coordinates and associated volumes for the cryo-EM reconstruction of GlyR-GABA-closed-SMA, GlyR-GABA-open-SMA, GlyR-GABA-desensitized-SMA and GlyR-GABA-super-open-SMA data sets have been deposited in the PDB and EMDB under the accession codes 6PLZ and 20381, 6PLY and EMD-20380, 6PLX and 20379, and 6PLW and 20378. The coordinates and associated volumes for the cryo-EM reconstruction of YGF-GABA-open-SMA, YGF-GABA-desensitized-SMA and YGF-GABA-super-open-SMA data sets have been deposited in the PDB and EMDB under the accession codes 6PLO and 20370, 6PLP and 20371, and 6PLQ and 20372. The coordinates and associated volume for apo-GlyR_EM_ data set have been deposited in the PDB and EMDB under the accession code 6PXD and 20518.

## Author Information Statement

The authors declare no competing interests. Correspondence and requests for materials should be addressed to Lucia Sivilotti or to Eric Gouaux

## Methods

### Protein purification and nanodisc reconstitution

The cDNA encoding the full-length zebrafish α1 glycine receptor (GlyR, NP_571477), with the following modifications, was cloned into the pFastBac1 vector for baculovirus expression in the Sf9 insect cells^20^. A thrombin cleavage sequence (Leu-Val-Pro-Arg-Ser) was introduced after the carboxy terminus of GlyR and before an octa-histidine tag. The truncation of GlyR (GlyR_EM_), in which the long M3-M4 loop replaced by an Ala-Gly-Thr tripeptide, were engineered based on the wild-type (WT) background. The YGF mutant, where residues Phe175 and Tyr177 were swapped, were generated using site-directed mutagenesis. WT receptor and YGF mutant were purified in the same procedures. Following transduction of insect cells with baculovirus and cell culture at 27 °C for 72 hours, cells were collected and resuspended in a buffer composed of 20 mM Tris pH 8.0 and 150 mM NaCl (TBS) in the presence of 0.8 µM aprotinin, 2 µg/ml leupeptin, 2 mM pepstain A and 1 mM phenylmethylsulfonyl fluoride and disrupted by sonication. Membranes were isolated by centrifugation for 1 h at 40,000 RPM (Ti45 rotor). The membrane fraction was collected and solubilized in 20 mM n-dodecyl-β-D-maltopyranoside (C12M) and 2.5 mM cholesteryl hemisuccinate Tris salt (CHS) for 2 hours with slow stirring at 4 °C. After ultracentrifugation to clarify the solution, the supernatant was collected and incubated with Talon resin for 1 hour and washed with TBS buffer supplemented with 1 mM C12M, 0.25 mM CHS and 35 mM imidazole. The receptor was eluted from the affinity resin using a buffer containing 250 mM imidazole. Fractions were pooled together and applied to a size-exclusion chromatography (SEC) Superose 6 10/300 GL column for further purification using a TBS buffer supplemented with 1 mM C12M and 0.25 mM CHS. The peak fractions were collected and concentrated to ∼16 µM receptor concentration for reconstitution into lipid nanodiscs or used for cryo-EM studies.

To remove unbound and bound glycine to enable determination of the apo-GlyR_EM_ structure, all the buffers were prepared using HPLC water (Sigma) and extensive dialysis was carried out as follows. The pelleted membrane with a volume of 30 mls was homogenized and incubated with 100 nM strychnine. The strychnine saturated membranes were transferred to an 8-10 kDa molecular weight cut off cellulose ester dialysis tubing (Spectra/PorBiotech) and dialyzed in 200-fold volume of glycine-free buffer containing 150 mM NaCl, 20 mM Tris 8.0 (TBS buffer in HPLC water) in the presence of 1 mM PMSF, 0.8 µM aprotinin, 2 µg/ml leupeptin, and 2 mM pepstatin A. Buffer exchange was carried out twice a day over the course of five days at 4°C, resulting in 10 changes.

To incorporate the receptor in lipid nanodiscs^51^, brain total lipid extract (BTL, Avanti) dissolved in chloroform was dried down using a rotary evaporator and resuspended in a TBS buffer supplemented with 20 mM C12M and 2.5 mM CHS to a final lipid concentration of 10 mM. Because we estimate that the diameter of the GlyR transmembrane domain is ∼80 Å, we used the MSP1E3D1 scaffold protein which, in turn, yields nanodiscs with an outer diameter of ∼12 nm. MSP1E3D1 was purified through metal chelate affinity column (Ni-NTA) and extensive dialysis against TBS buffer^52^. GlyR was mixed with MSP1E3D1 and BTL in a final ratio of 1: 7.5: 200 (molar ratios) and incubated for 40 min by gentle agitation at 4 °C. Detergent was removed by adding Bio-Beads (SM2, Bio-Rad) to a final concentration of 150 mg/ml. After incubation for 2 hours, the Bio-Beads were replaced with fresh beads. The mixture was incubated overnight at 4 °C. After separation from Bio-beads, the protein was loaded onto a Superose 6 10/300 GL column equilibrated with TBS buffer. Peak fractions corresponding to reconstituted GlyR were collected for cryo-EM analysis and scintillation-proximity assay (SPA) ^53^.

### Protein purification via styrene maleic acid (SMA) copolymer solubilization

The SMA copolymer XIRAN 30010 (∼2.3:1 molar ratio of stryrene: maleic acid) was purchased from Polyscope as an aqueous solution of 20% (w/v) SMA. The purification procedures in the context of the SMA copolymer are similar to those previously described except for the following differences. Following collection of the membrane fraction by centrifugation, the membranes were resuspended in TBS buffer and XIRAN 30010 was added to a final concentration of 0.5% with slow stirring at 4 °C for 1 h. After ultracentrifugation for 1 hr at 40,000 RPM (Ti45 rotor), the supernatant was collected and incubated with NTA metal ion affinity resin for 5-6 hours. The remaining purification steps were carried out as described above with the caveat that no additional SMA or detergents were added throughout the remaining purification steps. Following SEC chromatography, the WT/YGF mutant-SMA complex was concentrated to 2 mg/ml for cryo-EM analysis.

### Determination of glycine contamination

Potential glycine contamination in 50 mg/ml stocks of taurine or GABA was analyzed using liquid chromatography tandem mass spectrometry (LC-MS/MS) with electrospray ionization in positive mode. MRM (multiple reaction monitoring) transitions monitored specific product ion fragment from glycine. The reported major fragments were 75.9/48 and 75.9/30.2. Standards spiked with 0.1 µg/ml to 50 µg/ml glycine in presence of a constant concentration of 50 mg/ml of GABA or taurine were prepared as control to test assay sensitivity. The glycine contents in the 50 mg/ml stocks of taurine or GABA were below the lower limit of detection, 0.1 µg/ml. These experiments allowed us to estimate that the extent of glycine contamination was less than 1.33 µM of glycine in 0.4 M of taurine or 0.49 M of GABA.

### Ligand binding assay

Competition ligand binding experiments were carried out using the SPA ^53^ and a solution containing 40 nM WT receptor or YGF mutant reconstituted into the SMA, 1 mg/ml Cu-Ysi beads, 0.1% bovine serum albumin, TBS buffer and 100 nM ^3^H-labelled strychnine (1:9 ^3^H:^1^H). The stocks of cold competitors - glycine, taurine and GABA – were prepared in TBS at a final concentrations of 0.1M, 0.5M and 1M respectively, followed by serial dilution. To estimate the extent of non-specific binding, 250 mM imidazole was used for background subtraction. The binding reactions reached equilibration after an overnight (<12 hr) incubation and the scintillation counts from the beads were read on a Micro Beta TriLux 1450 LSC & Luminescence Counter (Perkin Elmer). Ki values were determined with the Cheng–Prusoff equation in GraphPad Prism.

### Cryo-EM sample preparation and data acquisition

To prepare the GlyR complexes with different agonists, the WT receptor complexes in nanodisc or SMA were mixed with 10 mM glycine, 20 mM taurine or 40 mM γ-aminobutyric acid (GABA) immediately before the samples were placed onto the grids. Apo-GlyR_EM_ sample was placed onto the grid directly. For preparing the YGF mutant complexes with 40 mM GABA, it followed the same procedures as WT receptor. We estimate that the time elapsed between mixing of receptor with agonist and plunge freezing of the grids was around 10 s. In addition, the extent of glycine contamination in the taurine and GABA chemicals was estimated as less than 1.33 µM in 0.4 M of taurine or 0.49 M of GABA, as described below. To minimize preferred orientation, the nanodisc-reconstituted receptor solutions were supplemented with 60 µM perfluoro octyl maltoside (FOM) prior to grid preparation. A 2.5 µL sample of receptor was applied to glow-discharged Quantifoil R1.2/1.3 or 2/2 200 mesh gold holey carbon grids which were then blotted for 2.5 s under 100% humidity at 12 °C. The grids were then flash-frozen in liquid ethane, maintained at melting temperature, using a FEI Mark IV cryo-plunge instrument.

Cryo-EM datasets for the receptor-nanodisc complexes were collected on a Talos Arctica microscope (FEI) operated at 200 kV. Images were acquired on a K2 Summit direct-detector (Gatan) using super-resolution mode at a magnification of x45,000, corresponding to a pixel size of 0.455 Å. The defocus range was set from −0.8 µm to −2.2 µm. Each micrograph was recorded over 100 frames at a dose rate of ∼5.49 e^−^/pixel/s and a total exposure time of 10 s, resulting in total dose rate of 61 e^−^/Å^2^.

For the wild type receptor complexes in SMA, all the datasets were collected on a 300 kV FEI Titan Krios microscope equipped with an energy filter (Gatan Image Filter) set to 20 eV. Images were acquired on a K2 Summit direct-detector positioned after the energy filter using super-resolution mode at 165,000 magnification and a pixel size of 0.412 Å. Images were collected using ‘multishot’ methods driven by SerialEM ^54^ with seven shots per hole utilizing 2/2 200 mesh grids and a defocus ranging from −1.0 µm to −2.0 µm. Each micrograph was recorded over 40 frames with a total dose of 42 e^−^/Å^2^. For the YGF mutant in complex with GABA, the dataset was collected on the same Titan Krios microscope as the wild type, but images were acquired on a K3 BioQuantum direct-detector using super-resolution mode at 135, 000 magnification with a pixel size of 0.65 Å. Each micrograph was recorded over 60 frames at a dose rate of ∼18.76 e^−^/pixel/s and a total exposure time of 1.5 s, resulting in total dose rate of 62 e^−^/Å^2^.

The data for apo-GlyR_EM_ was collected on a 300 kV FEI Titan Krios at the cryo-EM facility at the Janelia Research Campus. This microscope is equipped with a CETCOR Image Corrector for spherical aberration correction and an energy filter. A 20eV energy slit centered on the zero-loss peak, and a C_s_ of 0.01 mm were used during data collection. All micrographs were recorded on a Gatan K2 Summit direct electron detector operated in super-resolution counting mode with a pixel size of 1.04 Å. The dose-fractionated images were recorded using the automated acquisition program SerialEM^54^. Each micrograph was recorded over 30 frames with a total exposure time of 6 s, resulting in total dose rate of 54 e^−^/Å^2^.

### Image processing

Super-resolution image stacks were 2×2 binned in the Fourier space and motion corrected by Motion Cor2 ^55^. The contrast transfer function (CTF) parameters were estimated by Gctf ^56^ and particles were picked by DoG-picker ^57^ for all the datasets except for indicated.

We first analyzed the taurine-bound GlyR-nanodisc complex data, culling through the total micrographs and retaining 1,264 micrographs that had Thon rings extending to at least 4 Å. Approximately 669,253 particles were picked from these micrographs and subjected to 2D classification in Relion 2.1 ^58^. After several rounds of classification, 2D classes with clearly defined features of the glycine receptor contained 244,309 particles. This particle stack was then employed for subsequent analysis. An initial model generated from cryoSPARC ^59^ was then used for 3D classification and refinement. A soft mask with extending binary map of 4 pixels and a soft-edge of 6 pixels and C5 symmetry were applied during all the following steps of classification and refinement. Two of the 3D classes (class1 and class 3) showed reasonable secondary structure features and were subjected to 3D refinement, yielding two different maps belonging to what we define as the desensitized and closed states, respectively. To reduce heterogeneity among these two classes and improve the density in the transmembrane regions, class 1 and class 3 were re-classified into three classes. For class1, this resulted in an additional class 1a (45,300 particles) and class 1b (79,557 particles) with the same transmembrane helix orientation and density. We therefore combined these classes and carried out 3D refinement, which yielded a map that we identify as the taurine-bound desensitized state. For class 3, the resulting class 3b contained only 15,292 particles, albeit with a better density in the transmembrane region. This class was selected for the 3D refinement, yielding a map that we identify as the taurine-bound closed state (Extended Data Fig. 8 and Supplementary Fig. 1).

A similar procedure was used to process data associated with the GABA-bound receptor in nanodiscs. Briefly, 493,873 particles were auto-picked from 1,109 micrographs and were subjected to 2D and then 3D classification. Two different classes containing 39,926 and 38,383 particles were selected for 3D reconstruction and 3D auto-refinement respectively, giving rise to one map that we identify as the GABA-bound desensitized state and a second map that we believe represents the GABA-bound receptor in a closed state (Extended Data Fig. 9).

The single particle cryo-EM data for the glycine-bound receptor in nanodiscs was processed using similar procedures. To do this, 472,910 particles were auto-picked from 1,231 micrographs and subjected to 2D and 3D classification. Notably, the 3D classes shared the same overall shape and transmembrane helix orientation, suggesting the preponderance of one conformational state in this glycine-bound data set. Three classes (80,121 particles) with good secondary structure features were combined and used for 3D auto-refinement, yielding a map that we define as the glycine-bound desensitized state (Extended Data Fig. 7).

For the glycine bound receptor in the SMA, a total of 5309 micrographs were retained followed the same criteria as above. A total of 1,578k particles were picked by DoG-picker and Relion-3.0 ^60^ was used for 2D classification, 3D auto-refinement and 3D classification. 4× binned particles were extracted followed by one round of 2D classification by limiting the resolution to 8Å. After this round of 2D classification, only these classes with clear receptor features and clean background were selected. 804k ‘good’ particles were selected and 2× binned particles were re-extracted followed by one round of 3D refinement using a filtered map and an initial model derived from the previous structure determination of GlyR (PDB code: 3jaf). Additional 3D classification was performed using the refined 2× binned particles by applying C1 or C5 symmetry with alignment. The 3D classifications with C1 symmetry yielded similar classes as with C5 symmetry, albeit with slight weaker TM density, thus justifying the imposition of C5 symmetry. The 172k particles belong to the dominant class6 were selected and unbinned particles were re-extracted and re-centered followed by another round of 3D refinement. Using the refined unbinned particle stack, 6 classes were generated during the 3D classification without alignment. Based on the differences of the M2 helices in these classes, 84978, 155456 and 39586 particles belong to the putative open state (class 6a), the desensitized state (class 6b), and the ‘super-open’ state (class 6c) were re-extracted for 3D auto-refinement (Extended Data Fig. 7).

Similar procedures were performed for the taurine-bound SMA data set. Here, 5004 movies yielded 1124k particles that were automatically selected using template based particle picking in Relion-3.0^60^. 4× binned particles were extracted followed by one round of 2D classification. 543k ‘good’ particles were retained and unbinned particles were extracted. After 3D classification, four different classes containing 17873, 17271, 122322 and 74365 particles were isolated and were used in 3D auto-refinement, giving rise to the density maps of the taurine-bound closed, taurine-bound desensitized, taurine-bound open and taurine-bound super-open states, respectively (Extended Data Fig. 8 and Supplementary Fig. 2). For the GABA-bound complex in SMA, a total of 2062 k particles were extracted from 5,200 micrographs and were subjected to 2D and 3D classification that the classes without obvious receptor features were dumped. Four different classes containing 121470, 150199, 20845 and 31667 particles were selected for 3D reconstruction and 3D auto-refinement respectively, giving rise to four maps that we identify as super-open, open, desensitized and closed states, respectively (Extended Data Fig. 9). For the GABA-bound YGF mutant complex in SMA, a total of 1800k particles extracted from 9,300 micrographs were subjected to the following 2D and 3D classification. The resulted 407k particles produced two classes containing 32386, 27496 and 42097 particles that we identified as open, desensitized and super-open states, respectively (Extended Data Fig. 10).

For the apo-GlyR_EM_ in the micelle, approximately 1000 particles were manually picked for an initial reference-free 2D classification. Seven to eight representative 2D class averages were selected as templates for automated particle picking using Relion 2.1^58^ or GAUTOMATCH (www.mrc-lmb.cam.ac.uk/kzhang/Gautomatch/). The auto-picked particles were visually checked and false positives were removed, giving rise to 204, 323 total particles. The particles were further cleaned-up by 2D classification using Relion 2.1^58^. The reconstruction of the published gly-EM complex (EMD-6345) was low-pass filtered to 60 Å and used as an initial model. No symmetry was applied during 3D classification in Relion 2.1^58^. Classes with characteristic features of Cys loop receptors were then selected. Particles belonging to the chosen classes, with a number of 115, 864, were used for 3D auto-refinement (Supplementary Fig. 3).

To improve the density in the transmembrane region, all the refined maps from Relion 2.1/3.0^58, 60^ were further refined using auto refinement in cisTEM ^61^, yielding well-defined maps. The final reconstructions for the receptor in the nanodiscs were subjected to post-processing in the Relion 2.1, and for the reconstructions of the receptor in SMA, the maps were sharpened by Localscale^62^. Local resolution was determined by Relion 3.0^60^. The reported resolutions in the Extended Table 1 and Extended Table 2 were estimated by gold standard FSC 0.143 criteria^63^.

### Model building

The model building commenced with rigid body fitting of an initial model derived from the prior GlyR structures to each density map followed by real space refinement in Coot ^64^and by using Phenix ^65^. To build all of the desensitized protein structures, the taurine-bound desensitized state structure in nanodisc (taurine_desensitized_nanodisc) was first built and then was used as the initial structure for the other desensitized state complexes. We began by fitting the glycine/ivermectin-bound protein structure (PDB code: 3jaf), excluding the ligands, into the taurine-bound desensitized state map using UCSF Chimera ^66^. The resulting structure was manually adjusted in Coot, guided by well resolved side chain densities. Amino acid residues missing from the previously determined glycine/ivermectin protein structure were added, including H327 at the end of the M3 helix and E390, M391 and R392at the beginning of the M4 helix. Prominent tube-shaped electron density surrounding the receptor transmembrane domain (TMD) likely arises from ordered lipid molecules and thus alkane chains with appropriate lengths were placed in the density features. The structure was further refined using Phenix in the context of restraints for stereochemistry of the alkane chains and carbohydrate groups. After Phenix refinement, the map to model cross correlation (cc) value between the final model and the map was 0.83. To build the other desensitized state structures, the above determined structure taurine_desensitized_nanodisc was fitted to the density maps using UCSF chimera. After rigid-body fitting of the taurine_desensitized_nanodisc model, taurine was replaced with the appropriate ligands, alkane chains were either replaced with appropriate lengths in these nanodisc structures or removed in the SMA structures, and the models were then real-space refined in Coot and Phenix. Specifically, two amino acids - V388 and E399 - were added at the beginning of the M4 helix in these protein structures of the desensitized state in SMA.

The procedures to the build open/super-open state protein structures were similar to the model building of the desensitized protein structures. The glycine-bound open state protein structure in SMA (glycine_open_SMA) was achieved by fitting the glycine-bound desensitized protein structure in SMA (glycine_desensitized_SMA) to its corresponding map using UCSF Chimera. Coot was then used to manually adjust the glycine_desensitized_SMA structure, mainly focusing on the TMD, followed by Phenix refinement, yielding a map to model cc of 0.80. To build the taurine-bound and GABA-bound open/super-open state protein structures derived from the receptor-SMA complex, the glycine_open_SMA structure was simply fit to the corresponding maps using UCSF Chimera, following by fitting of the appropriate ligands to the density in the neurotransmitter binding pocket. Phenix real space refinement was next performed to refine the protein structures within their corresponding density maps.

To build the GABA-bound YGF mutant open, desensitized and super-open states, the two amino acids 175F and 177Y were firstly mutated into 175Y and 177F by coot, respectively, within their corresponding wild type GABA-bound structures. After Phenix refinement, the resulted map to model cc values were 0.80, 0.79 and 0.72 for YGF mutant open, desensitized and super-open states, respectively.

For model building of the taurine-bound closed state in nanodisc (taurine_closed_nanodisc), we found that the previously determined strychnine-bound structure (PDB code: 3jad) fit the map well and thus it was used as an initial model. The structure was fit to the taurine-bound closed state map using UCSF Chimera. Chain A of the superimposed model was further optimized by real space refinement using Coot, including fitting of a taurine molecule to the density in the neurotransmitter binding pocket, adding Q326 and H327 at the end of the M3 and E390, M391 and R392 at the beginning of M4 helices, manually building the M2-M3 loop and adding lipid-derived alkane chains surrounding TMD. The model generated in Coot was further refined against the density map by Phenix. The cc between the taurine_closed_nanodisc and its corresponding map was 0.81. To build the other state protein structures either in nanodisc or SMA, the Phenix refined taurine_closed_nanodisc coordinates were used as the initial structure. Coot was used to change the ligands and removed the alkane chains in SMA structures or replaced with appropriate lengths in nanodisc structures based on the density features, followed by Phenix refinement. All of the final models have good stereochemistry as evaluated by molprobity^67^ (Supplementary Tables 1-4). All the figures were prepared by Pymol (Schrödinger, LLC) and UCSF chimera^68^. Pore radii were calculated using program Hole^69^.

### Molecular dynamics simulations

Two types of molecular dynamics (MD) simulations were conducted to investigate the permeation properties of the experimentally derived structural models, respectively. The membrane composition used in the simulations corresponded to the HEK cytoplasmic membrane (24% POPS, 7% POPI, 3% POPG, 11% sphingomyelin, 16% POPE, 36% POPC, and 3% cholesterol). Only the transmembrane domain (residue 244-436) was simulated for the glycine-bound desensitized, open, and super-open state structural models embedded in lipid bilayers for 500 ns each. To verify that these channel structures can be blocked by toxins as observed experimentally, an additional set of simulations were performed in the presence of a channel blocker, picrotoxinin, for each of the three states using the same protocol for 500 ns.

CHARMM-GUI^70^ was used to construct the initial lipid bilayers with a cross-sectional of 180 × 180 Å^2^. VMD^71^ was used for other steps of system preparation. The proteins were embedded into the membranes as previously described^25^. A picrotoxinin was added to each protein near the −2’P gate when preparing for ion channel blocking studies. Then, all systems were neutralized and solvated in a 150 mM NaCl. Each system had a final dimension of 180 × 180 × 150 Å^3^. NAMD 2.12^72^ was used to perform all of the simulations.

Each simulation started with 3,000 steps of energy minimization with phospholipid phosphorus atoms, hydroxyl oxygen of cholesterol, and protein side chain heavy atoms restrained (k = 1 kcal.mol^-1^.Å^-2^), along with additional restraints on the protein backbone atoms (k = 50 kcal.mol^-1^.Å^-2^), followed by a 0.5 ns equilibration of lipid tails. To relax the membrane lipids around the protein, a 20-ns simulation was performed where only protein backbone and ligand heavy atoms were restrained (k = 50 kcal.mol^-1^.Å^-2^). Then, the simulations continued with only protein backbone atoms restrained (k = 1 kcal.mol^-1^.Å^-2^), and with a transmembrane voltage of −100 mV applied throughout.

To maintain the constant pressure of 1 atm in our NPT simulations, the Nosé-Hoover piston^73, 74^ with an oscillation period of 200 fs and a damping time scale of 100 fs was used. A constant temperature of 310 K was maintained by coupling all of the non-hydrogen atoms to a Langevin thermostat with a damping coefficient of 1.0 ps^-1^. The SETTLE algorithm^75^ was used to keep the covalent bonds involving hydrogens rigid. The TIP3P^76^ model was used for the water molecules. We chose 2 fs time steps for the simulations. To calculate the long-range electrostatic interactions, the PME method^77, 78^ with a grid density of 1 Å^-3^ was used. For van der Waals and short-range electrostatic interactions, we used a smooth cutoff of 12 Å. CHARMM36m^79^ force field parameters were used for the protein, and CHARMM36 force field parameters^80^ for the lipids and ions. The parameters for picrotoxin were obtained from CGenFF^81, 82^. VMD was used for analyzing the MD trajectories. To calculate the pore-radius profiles, we used HOLE^69^.

### Patch-clamp and single-channel recording

#### Heterologous expression in human embryonic kidney cells

Full-length zebrafish α1 GlyR was subcloned into the pcDNA3.1 for expression in the human embryonic kidney 293A cells (HEK293A). Point mutations were using site-directed mutagenesis. HEK293A cells were transiently transfected using the calcium phosphate-DNA precipitation method. The DNA mix consisted of pcDNA3.1 plasmids with inserts coding for the enhanced Green Fluorescence Protein (eGFP; GenBank accession number U55763), and GlyR (wild-type or mutant). Plasmid without an open reading frame (“empty”) was added to the transfection mix to optimize the expression level^83^. The final mixture of cDNA contained 2% α1, 18% eEGFP and 80% empty pcDNA3.1 plasmid. Cells were washed 5-16 hrs later and electrophysiological experiments were performed 1-2 days after transfection.

#### Patch-clamp recordings and analysis

Macroscopic and single-channel currents were recorded from transfected HEK293A cells in the whole-cell, outside-out and cell-attached patch-clamp configurations at 20°C. Patch electrodes were pulled from thick-walled borosilicate glass capillary tubes (GC150F-7.5; Harvard Apparatus) on a Flaming/Brown puller (Model-P97, Sutter Instruments) and were fire-polished to give a final resistance of 3-5 MΩ for whole-cell recordings and 5-15 MΩ for cell-attached or outside-out recordings (when filled with the appropriate internal solution).

Cells were bathed in an extracellular solution consisting of (in mM): 20 Na D-gluconate, 112.7 NaCl, 2 KCl, 2 CaCl2, 1.2 MgCl2, 10 HEPES (4-(2-hydroxyethyl)-1-piperazineethanesulfonic acid), 40 glucose (all Sigma) and 10 tetraethylammonium chloride (TEACl) (Alfa Aesar); the pH adjusted to 7.4 with NaOH. For the cell-attached recordings, pipettes were filled with extracellular solution containing the required concentration of agonist. To record macroscopic currents, the pipette was filled with a 30 mM chloride intracellular solution, containing (in mM): 101.1 K gluconate, 1 ethylene glycol tetraacetic acid (EGTA), 1 CaCl2, 1 MgCl2, 10 HEPES, 2 MgATP, 40 sucrose, 6 KCl (all Sigma) and 20 TEACl, the pH adjusted to 7.4 with NaOH. All solutions were prepared in bi-distilled water or, for cell-attached single channel recordings, in high-performance liquid chromatography grade (HPLC) water (VWR Chemicals), and filtered through a 0.2 µm cellulose nitrate membrane filter (Whatman) before use. Glycine, taurine or GABA-containing solutions were prepared by diluting a 1 M glycine, 1 M taurine or 1 M GABA stock in extracellular solution.

Taurine and GABA were purified by 2x recrystallization from aqueous ethanol and subsequently tested for glycine contamination by an HPLC assay: samples were resuspended in 10 µl 50% EtOH, reacted for 30 min with 90% EtOH:triethylamine:phenylisothiocyanate (7:2:1), evaporated to dryness at room temperature and redissolved in 100 µl 5% acetonitrile in 0.1 M ammonium acetate. A 150×4.6 mm Hypersil C18 column was used and were detected was measured at 254 nm. The purified taurine and GABA contained no detectable glycine, which in this assay is less than 1 part in 10,000.

#### Whole-cell recording

Whole-cell currents were recorded from isolated transfected cells held at a holding potential of −40 mV (corrected for the junction potential of 11 mV) with an Axopatch 200B (Molecular Devices) and filtered at 5 kHz by the amplifier’s low pass 4-pole Bessel filter. Recordings were digitized with a Digidata 1440A (Molecular Devices) at a sampling frequency of 20 kHz and acquired to PC using Clampex 10.2 software (Molecular Devices) for offline analysis.

Agonist solutions were applied to the cells for approximately 1-2 s by a U-tube application system^84^, with a 10-90% exchange time <1 ms (as tested by application of 50% diluted extracellular solution to the open tip pipette). The saturating agonist concentration was applied every third or fourth application to check response stability throughout the recording. Only cells where the test response rundown was less than 30% during the experiment were accepted for further analysis. No correction for rundown was applied. Different agonist concentrations were applied in random order to obtain concentration-response curves. The peak current amplitudes for each glycine application were measured using Clampfit 10.2 software (Molecular Devices). The concentration-response data for each cell were then fitted with the Hill equation using the CVFIT programme (DCPROGS; https://github.com/DCPROGS/CVFIT) to estimate EC50 (agonist concentration required to elicit 50% maximal response), nH¬ (Hill-slope) and Imax (maximum peak current).

#### Fast agonist application

Macroscopic currents evoked in outside-out patches by fast agonist application pulses were recorded at a pipette holding potential of −100 mV (−111 mV when corrected for a junction potential of 11 mV). The internal solution was the same as the one for whole cell recordings and contained 30 mM chloride. Agonist, dissolved in extracellular solution, was applied to outside-out patches with a theta tube (Hilgenberg GmbH) cut to a final diameter of ≈ 150 μm at the tip. The tube was driven by a piezo stepper PZ-150M (Burleigh Instruments Inc). The exchange time was measured by the application of diluted bath solution (eg 30:70 bath solution: water) before the experiment (to optimize the electrode position) and after the rupture of the patch. The rise and decay times for these open-tip currents were measured using Clampfit 10.2 as the times from 20 to 80% of the peak response. Patches in which the open tip response had a 20–80% exchange time slower than 200 μs were rejected from further analysis. In order to study the kinetics of macroscopic currents, 10–20 responses were recorded in response to pulses of agonist applied at intervals of at least 10 s and averaged. Only experiments in which the rundown between the first and last three responses was < 30% were included in the analysis. The time course of the macroscopic currents was characterized by fitting the rise time between 20 and 100% and the decay time from 80 to 20% of the peak response with one or more exponentials.

#### Single-channel recording

Low-noise single-channel currents were recorded in the cell-attached configuration at a pipette holding potential of +100 mV with an Axopatch 200B and filtered at 10 kHz using the amplifier’s low pass 4-pole Bessel filter. The data was digitized with a Digidata 1322A (Molecular Devices) at a sampling frequency of 100 kHz and acquired to PC using Clampex 10.2 for offline analysis.

#### Single channel current amplitude and cluster *P*_open_ measurements

In order to compare single-channel current amplitudes and cluster open probability, single-channel recordings were filtered offline using the Clampfit 10.2 low-pass Gaussian filter with a final cut-off of 5 kHz and resampled at 50 kHz. At high agonist concentration, channel openings occurred in clusters delimited by long closed intervals, likely to be desensitized. These clusters are likely to originate from the activity of a single ion channel molecule and were used for *P*_open_ measurements. Clusters longer than 100 ms with more than 10 openings were selected for analysis. The gap between clusters was at least 100 ms or 300 ms (for GABA measurements). Channel activity in the selected clusters was idealized using the half-amplitude threshold method implemented in Clampfit 10.2 and open probability was calculated as the ratio of cluster open time over total cluster length. The amplitude of single channel currents was measured in Clampfit 10.2 as the average of all detected opening amplitudes inside a cluster.

## Extended Data Figure Legends

**Extended Data Figure 1**

**a,** Macroscopic responses of GlyR to the rapid application of 10 mM glycine, 100 mM taurine and 100 mM GABA to the same outside-out patch.

**b-c,** Ion permeation pathways and pore radius plot for the desensitized state of GlyR in complex with glycine in the SMA.

**d-e,** Conformational changes in the M2 helices between the desensitized and closed states of GlyR complexed with taurine (d) and GABA (e) in the nanodisc. Two constriction sites 9’L and - 2’P are shown as sticks.

**f,** Plots for pore radius as a function of distance along the pore axis for desensitized and closed states of GlyR complexed with taurine and GABA in the nanodisc as well as the apo-GlyR_EM_ in the micelle.

**Extended Data Figure 2**

**a,** A single subunit of glycine-bound open state of GlyR in the SMA viewed parallel to the membrane with secondary structure element denoted.

**b,** The minimum pore-radius of the open, desensitized, and super-open models of glycine-bound GlyR in the SMA along the 500-ns ion-permeation MD simulations with (left) or without (right) a picrotoxinin molecules placed in the pore.

**Extended Data Figure 3 Summary of Cα distances between two constriction sites 9’L or - 2’P of all the structures solved in the SMA for wildtype receptor and YGF mutant.**

The five M2 helices are isolated and shown in cartoon. The −2’Pro and 9’Leu are shown in sticks. The Cα distances between two −2’P or 9’L from adjacent subunits are shown in angstrom.

**Extended Data Figure 4**

Cryo-EM densities and models for the pore lining M2 helices in the super-open states of GlyR or YGF mutant complexes solved in the SMA to show the extra densities in the intracellular end of M2 helices. The constriction site −2’Pro is indicated.

**Extended Data Figure 5 The differences in the binding pocket between GlyR and α1β1γ2 GABA receptor (GABAR, PDB code: 6DW1) when bound with GABA**

**a,** Reference model for extracellular domain (ECD) of open state of GlyR complex with GABA in the SMA.

**b,** The enlarged view of the differences in the binding pocket after superposing the ECDs. The discrepancies residues are highlighted with rectangle. The Cα atoms of key residues are represented as spheres.

**c,** Schematic diagram illustrating the changes in distances of the Cα atoms of R81, F175, S145 and T220 in the GlyR and GABAR.

**d,** The open probabilities (P_open_) of wildtype (WT) and triple mutant (F115Y+F175Y+F223Y) of GlyR elicited by 100 mM GABA.

**Extended Data Figure 6**

**a-e**, Superposition of two adjacent subunits from the closed state of GlyR in complex with taurine in the SMA (tau-closed) with resting state of GlyR (apo-GlyR_EM_) and 5-HT3A receptor (5-HT_3A_R) (PDB code: 6BE1) to show the differences in the binding pocket (black box) and ECD-TMD interfaces (red box). The tau-closed, apo-GlyR_EM_ and 5-HT_3A_R are colored in cyan, red and purple, respectively. Taurine is shown in sticks.

**f-g**, Conformational changes in the TMD between tau-closed, tau-open and tau-desensitized states. The centers of mass (COM) of the four TM helices from one subunit is indicated. The rotation angle of the COM relative to the receptor center is labeled. The tau-closed, tau-open and tau-desensitized are in cyan, yellow and salmon, respectively.

**Extended Data Figure 7 3D reconstructions for glycine-bound states in the nanodisc (green box) and SMA (blue box).**

**a**, SEC trace for GlyR in the nanodisc and the SDS-PAGE for the peak fractions.

**b**, A typical cryo-EM micrograph for glycine-bound GlyR in the nanodisc with a 30 nm scale bar.

**c**, Selected 2D class averages for glycine-bound GlyR in the nanodisc.

**d**, Local resolution map for unsharpened glycine-bound GlyR in the nanodisc.

**e**, FSC curves for glycine-bound GlyR map before (unmasked) and after (masked) Relion’s post processing, and between the model and the final map.

**f**, Particle angular distribution for glycine-bound GlyR in the nanodisc.

**g**, SEC trace for glycine-bound GlyR in SMA with the SDS-PAGE for the peak fraction.

**h**, A typical cryo-EM micrograph for glycine-bound GlyR in the SMA with a 30 nm scale bar.

**i**, Selected 2D class averages for glycine-bound GlyR in the SMA.

**j**, **m and p**, Local resolution maps for unsharpened open, desensitized and super-open maps, respectively.

**k**, **n and q**, FSC curves before (unmasked) and after (masked) Relion’s post processing, and between the model and the final maps for open, desensitized and super-open, respectively.

**l**, **o** and **r**, Particle angular distributions for open, desensitized and super-open maps, respectively.

**Extended Data Figure 8 3D reconstructions for taurine-bound states in the nanodisc (green box) and SMA (blue box).**

**a**, A typical cryo-EM micrograph for taurine-bound GlyR in the nanodisc with a 30 nm scale bar.

**b**, Selected 2D class averages for taurine-bound GlyR in the nanodisc.

**c** and **e**, Local resolution maps for unsharpened taurine bound desensitized and closed states in the nanodisc, respectively.

**d** and **f**, FSC curves before (unmasked) and after (masked) Relion’s post processing, and between the model and the final map for taurine bound GlyR at desensitized and closed states, respectively.

**g**, A typical cryo-EM micrograph for taurine bound GlyR in the SMA with a 30 nm scale bar.

**h**, Selected 2D class averages for taurine bound GlyR in the SMA.

**i**, **k**, **m**, and **o**, Local resolution maps for unsharpened open, desensitized, closed and super-open, respectively.

**j**, **l**, **n** and **p**, FSC curves before (unmasked) and after (masked) Relion’s post processing, and between the model and the final map for open, desensitized, closed and super-open, respectively.

**Extended Data Figure 9 3D reconstructions for GABA-bound states in the nanodisc (green box) and SMA (blue box).**

**a**, A typical cryo-EM micrograph for GABA-bound GlyR in the nanodisc with a 30 nm scale bar.

**b**, Selected 2D class averages for GABA-bound GlyR in the nanodisc.

**c** and **e**, Local resolution maps for unsharpened GABA-bound desensitized and closed states in nanodisc, respectively.

**d** and **f**, FSC curves before (unmasked) and after (masked) Relion’s post processing, and between the model and the final map for GABA-bound GlyR at desensitized and closed states, respectively.

**g**, A typical cryoEM micrograph for taurine-bound GlyR in the SMA with a 30 nm scale bar.

**h**, Selected 2D class averages for taurine-bound GlyR in the SMA.

**i**, **k**, **m**, and **o**, Local resolution maps for unsharpened open, desensitized, closed and super-open, respectively.

**j**, **l**, **n** and **p**, FSC curves before (unmasked) and after (masked) Relion’s post processing, and between the model and the final map for open, desensitized, closed and super-open, respectively.

**Extended Data Figure 10 3D reconstructions for GABA-bound states of YGF mutant in the SMA**

**a**, A typical cryo-EM micrograph with a 30 nm scale bar.

**b**, Selected 2D class averages.

**c, f,** and **i,** Local resolution maps for unsharpened open, desensitized and super-open states.

**d, g** and **j**, FSC curves before (unmasked) and after (masked) Relion’s post processing, and between the model and the final map for open, desensitized and super-open states, respectively.

**e**, **h** and **k**, Particle angular distributions for open, desensitized and super-open maps, respectively.

## Supplementary Data

**Supplementary Figure 1**

Data processing flow chart for taurine-bound GlyR in the nanodisc.

**Supplementary Figure 2**

Data processing flow chart for taurine-bound GlyR in the SMA.

**Supplementary Figure 3 3D reconstructions for apo-GlyR_EM_ in the micelle**

**a**, A typical cryo-EM micrograph with a 20 nm scale bar.

**b**, Selected 2D class averages.

**c, d, e** Local resolution map, FSC curve and Particle angular distributions for apo-GlyR_EM_.

**Supplementary Figure 4 Representative densities of glycine or taurine bound maps**

The structures are shown in stick and maps are in mesh. For each panel, M1 and M2 helices together with M1-M2 loop starting from Y239 to S286 are isolated, contoured at 6.5 σ to 8 σ. One of the alkane chains is shown in the nanodisc maps. The M2-M3 loop from A288 to Y295 is illustrated, contoured at 6.5 σ to 8 σ. The β2 sheet from ECD from D73 to W84 is isolated, contoured at 7.5 σ to 8 σ. The nomenclature here is following the rule Ligand_State_MembraneMimic. The maps in nanodisc and SMA are sharpened by Relion and Localscale, respectively.

**Supplementary Figure 5 Representative densities of GABA bound maps**

Similar to Supplementary Figure 4, the related densities are shown for GABA bound maps in a same style, either in nanodisc and SMA. The nomenclature is also following the rule Ligand_State_MembraneMimic.

**Supplementary Figure 6 Partial agonists (taurine/GABA) densities in the closed states of GlyR in the nanodisc and SMA.**

GABA densities in the nanodisc and SMA are contoured at 3 σ and 4 σ, respectively. Taurine densities in the nanodisc and SMA are contoured at 4 σ and 0.02 σ, respectively.

**Supplementary Table 1**

Statistics for 3D reconstruction and model refinement in the nanodisc.

**Supplementary Table 2**

Statistics for 3D reconstruction and model refinement of glycine and taurine bound GlyR in the SMA.

**Supplementary Table 3**

Statistics for 3D reconstruction and model refinement of the GABA bound GlyR in the SMA.

**Supplementary Table 4**

Statistics for 3D reconstruction and model refinement of YGF mutant and apo-GlyR_EM_ in the micelle.

**Supplementary Table 5**

Resolution summary for all the maps.

**Supplementary Table 6**

Binding pocket metrics for Cys-loop family.

