## Supplementary figures and images for "Mechanism of gating and partial agonist action in the glycine receptor"

### Extended Figures

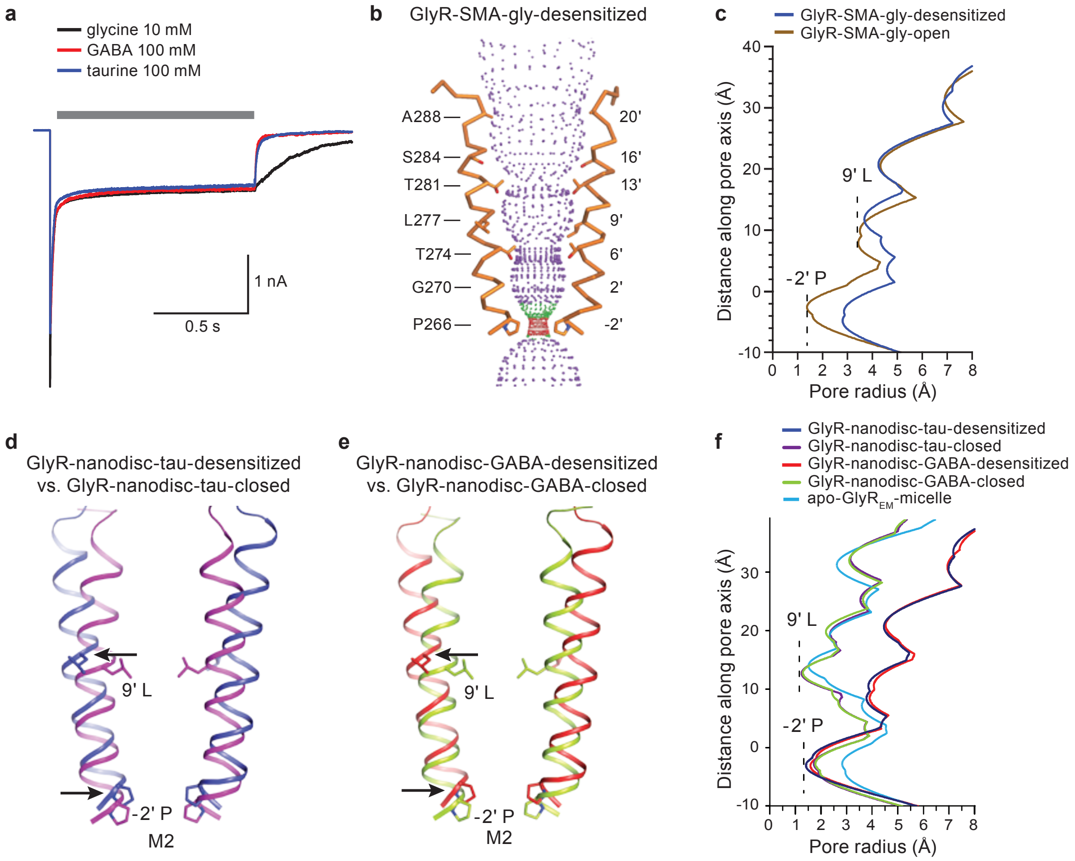

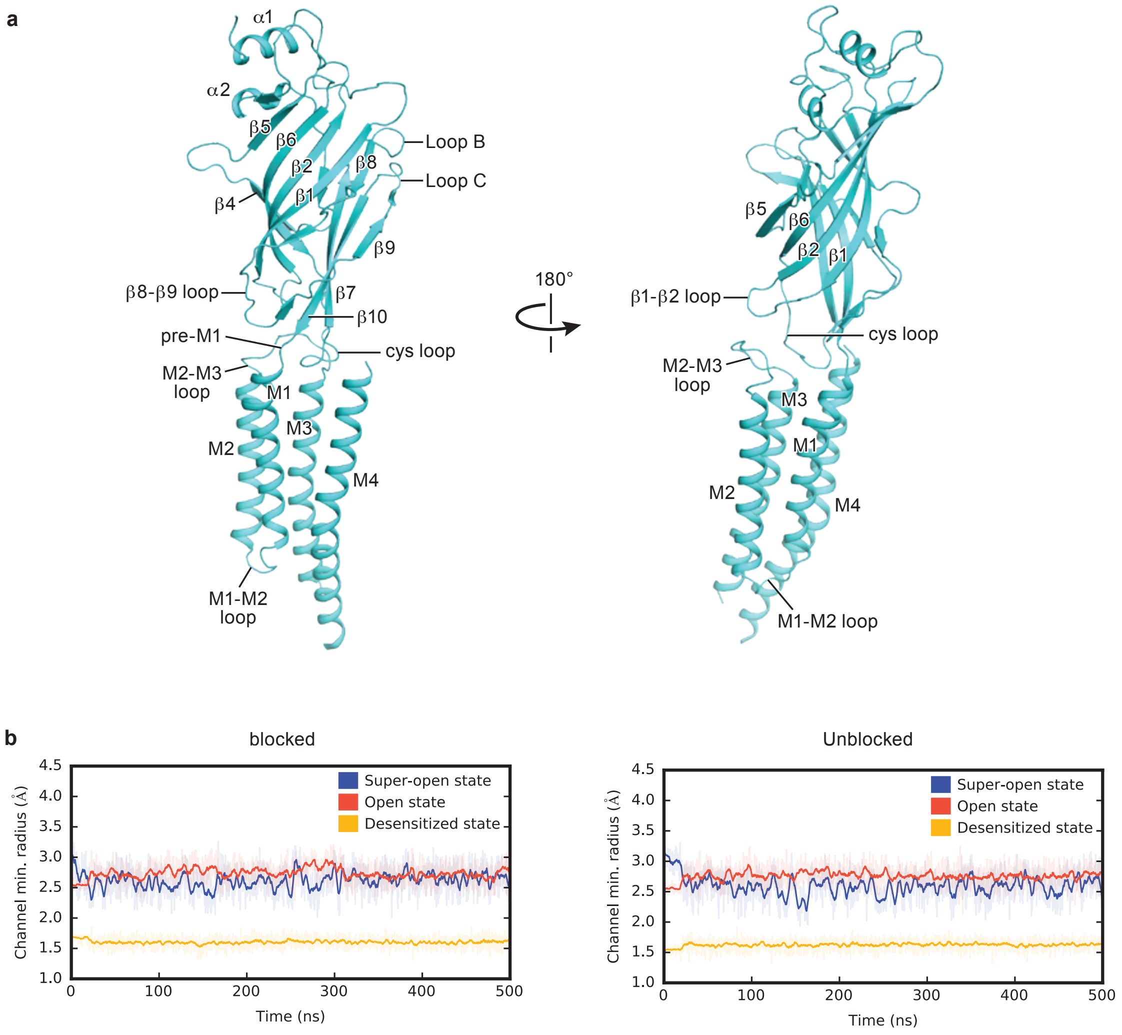

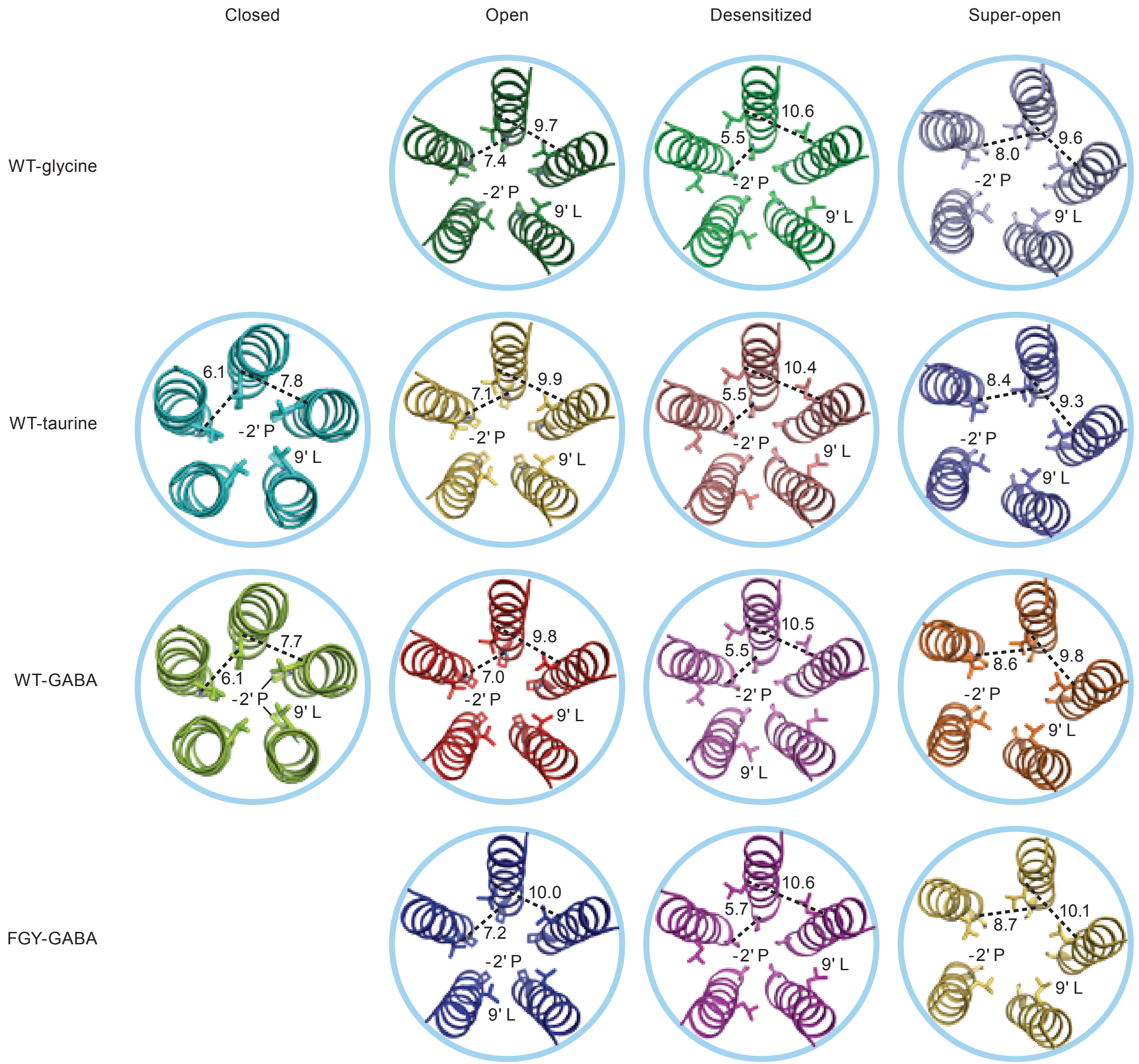

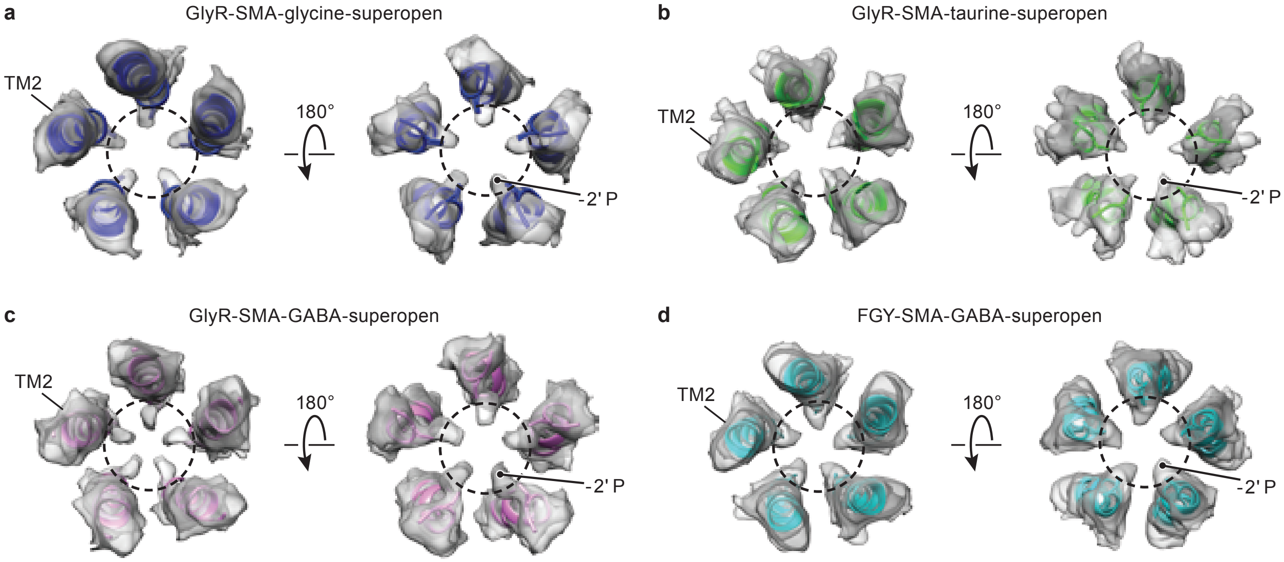

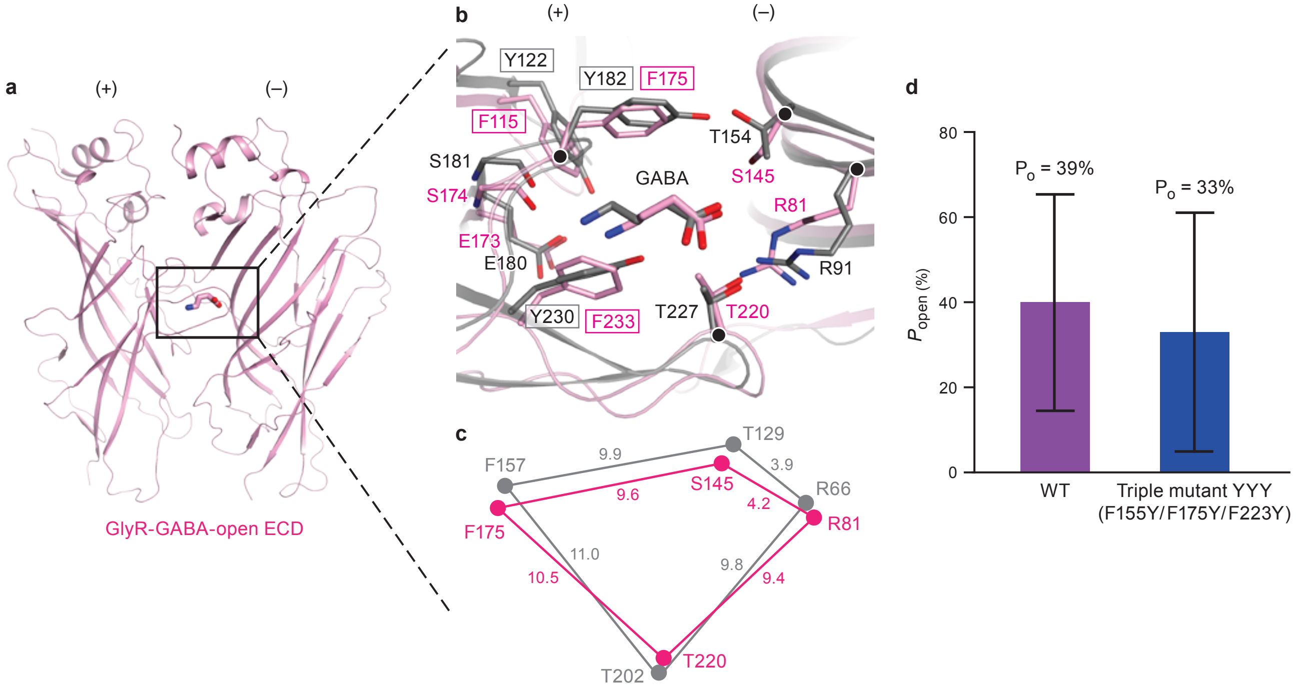

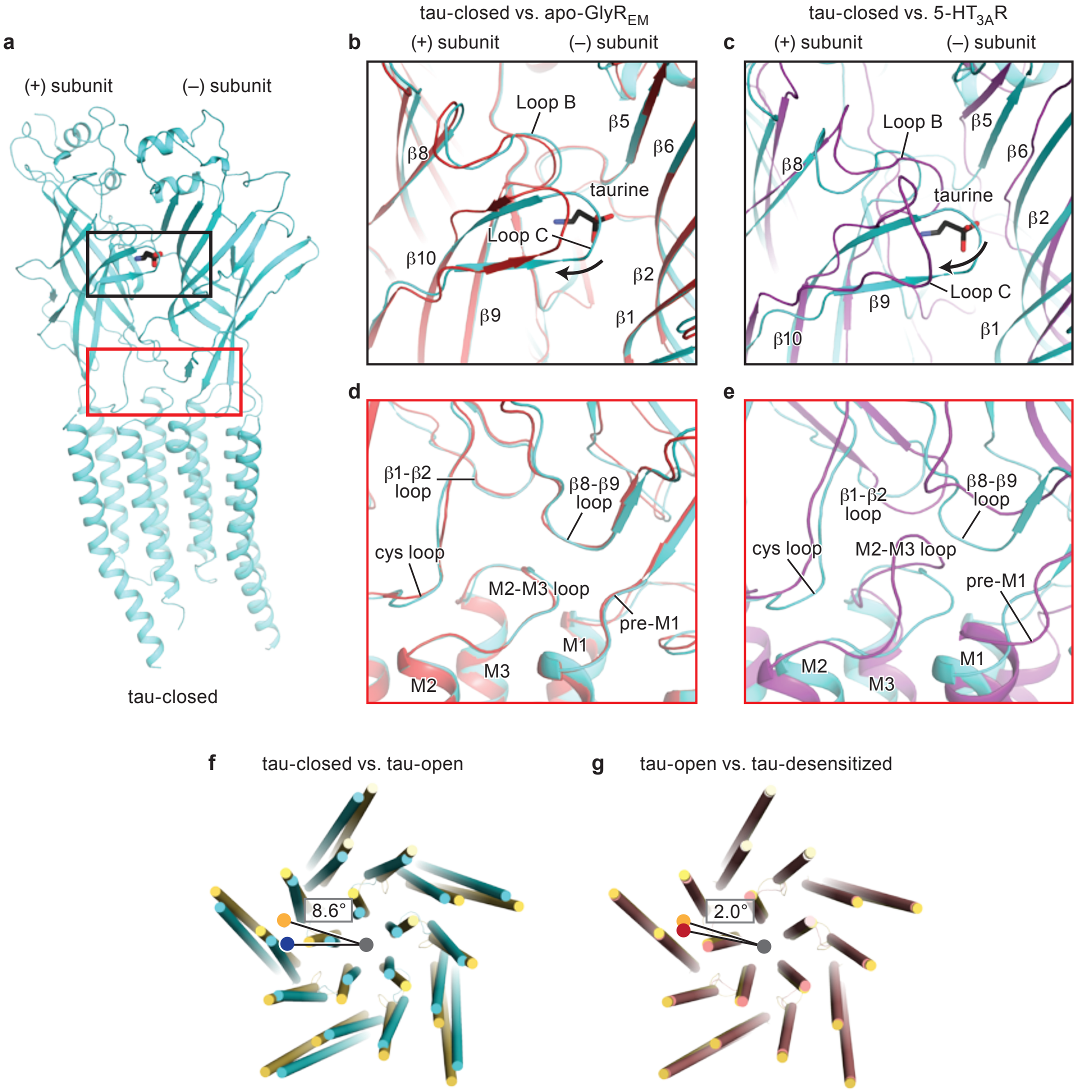

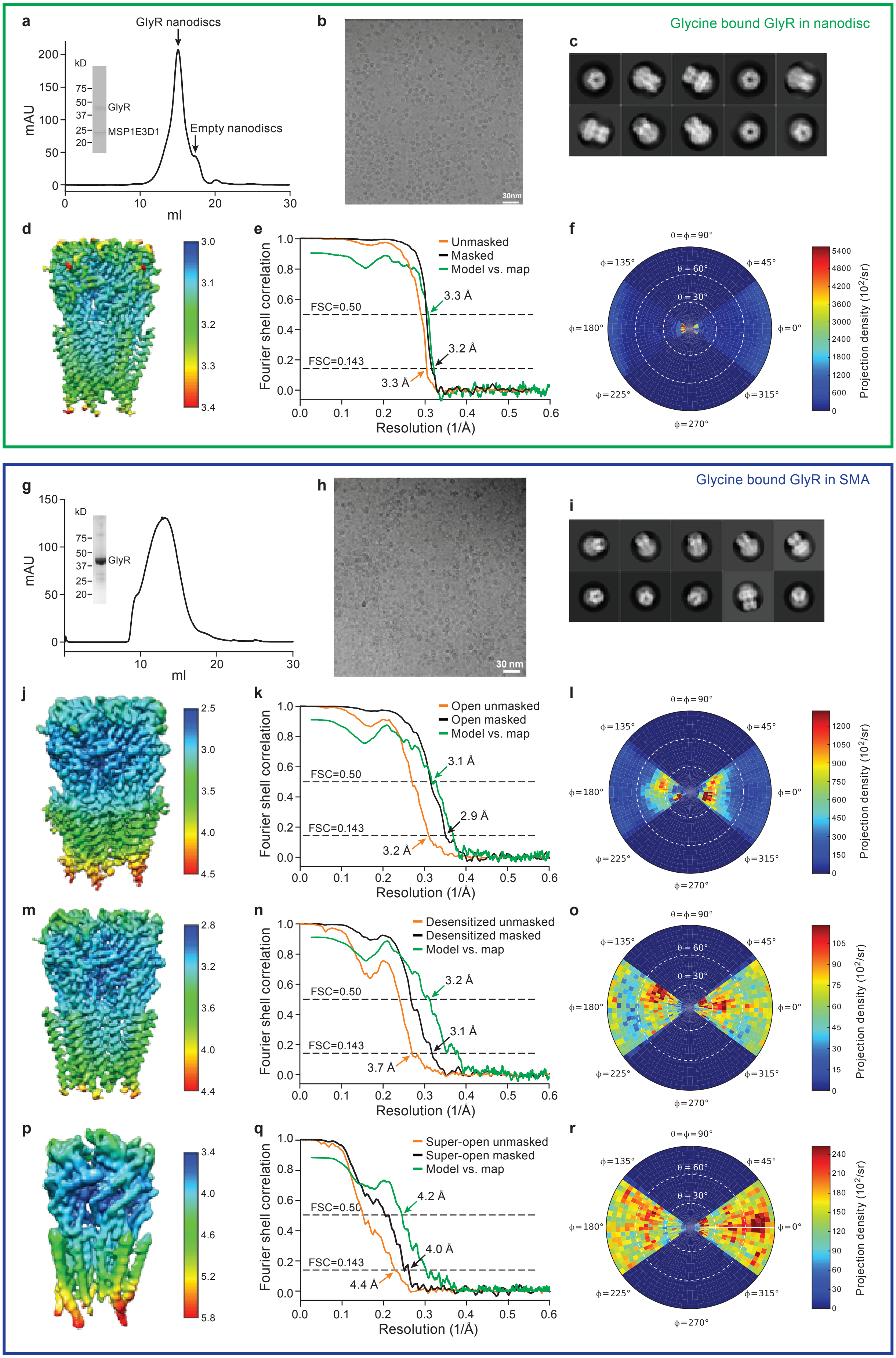

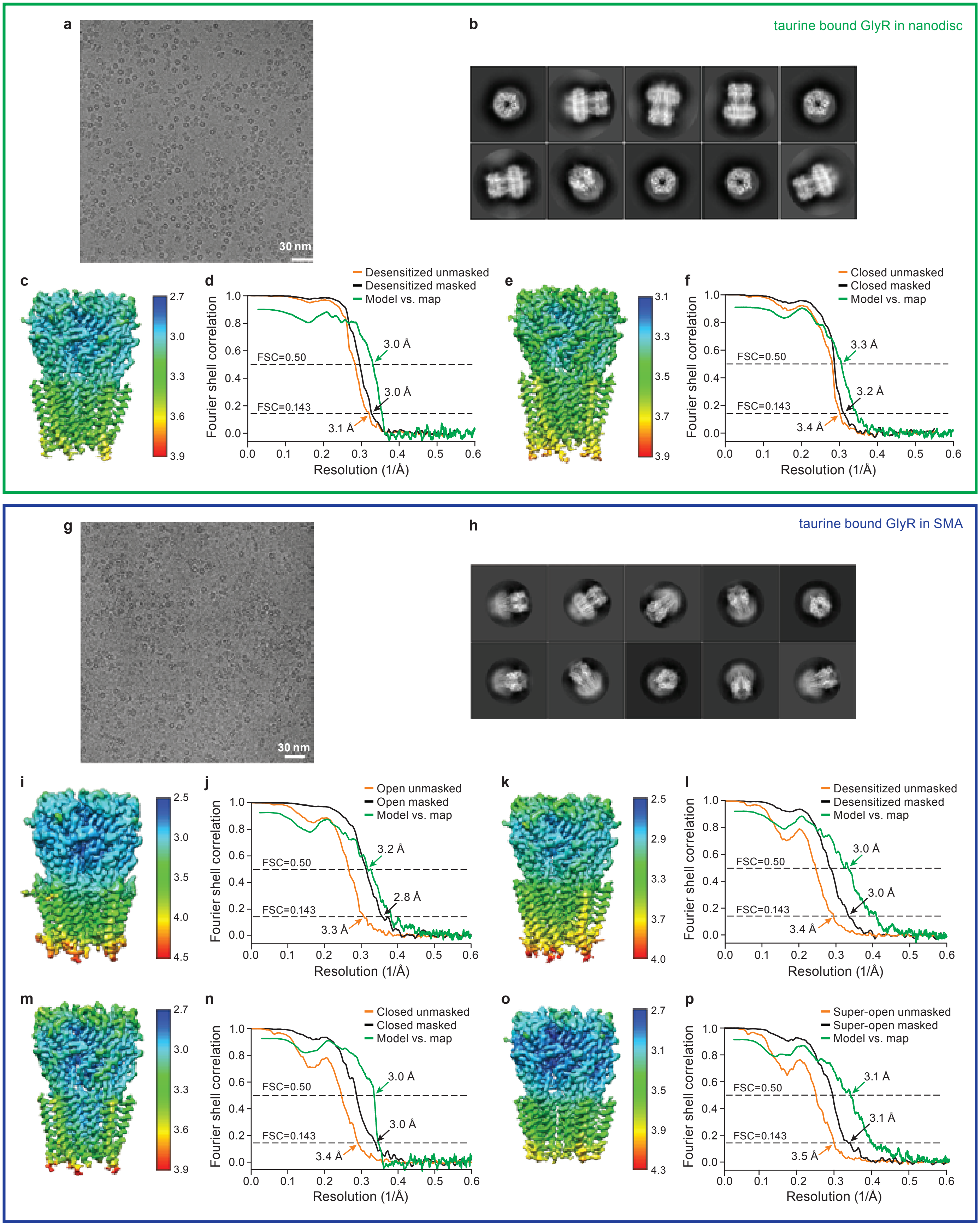

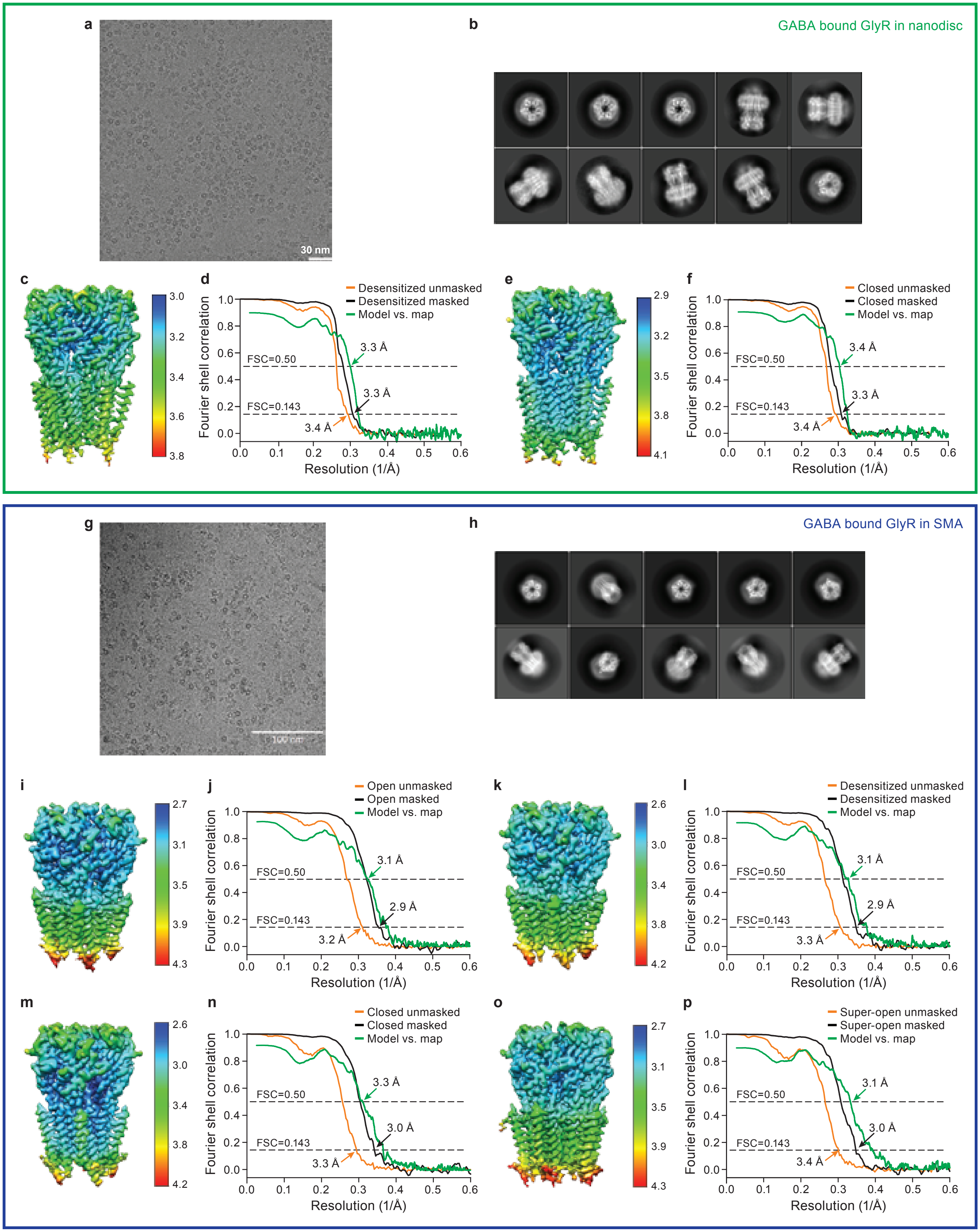

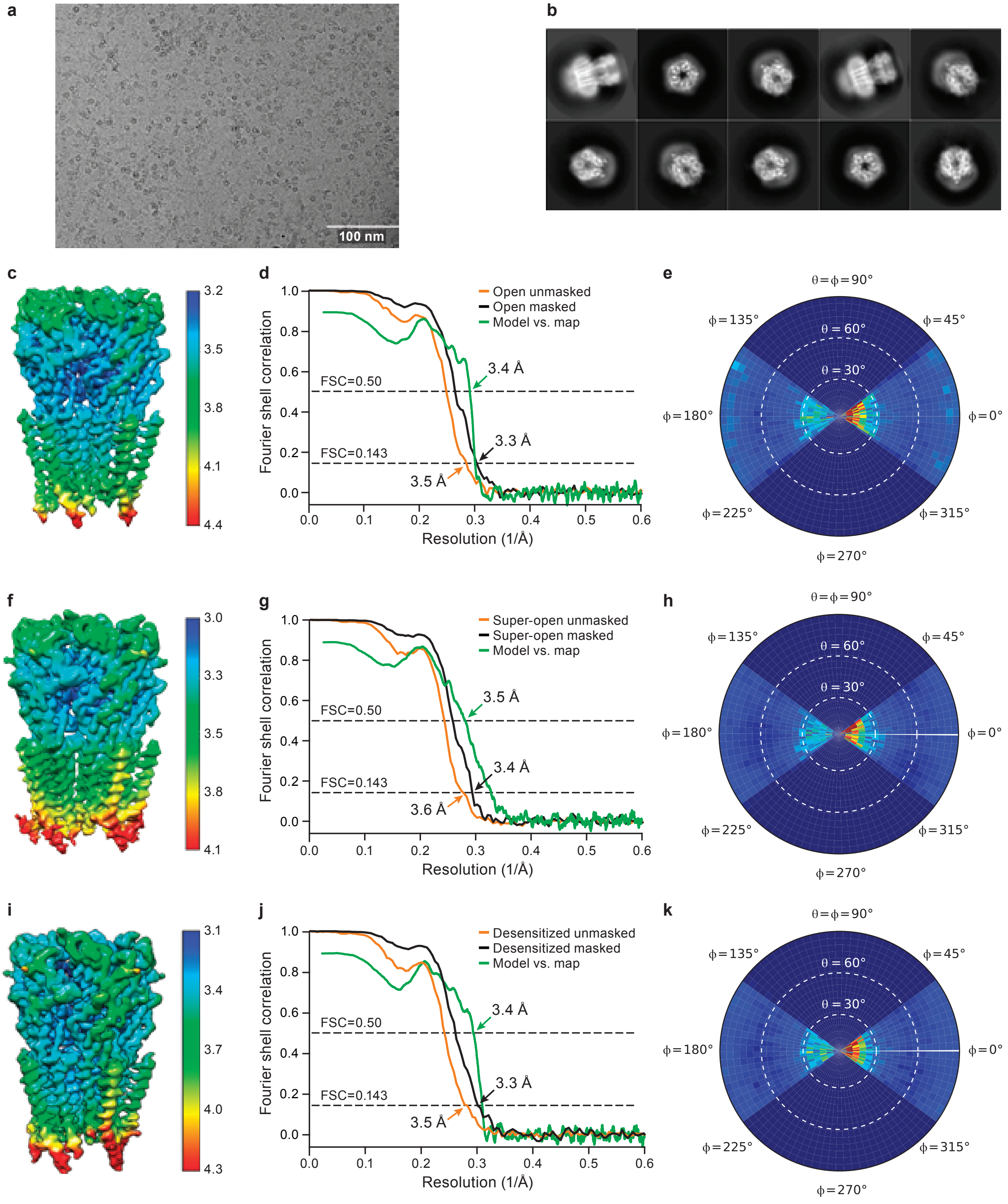
