## Supplementary Data for "Mechanism of gating and partial agonist action in the glycine receptor"

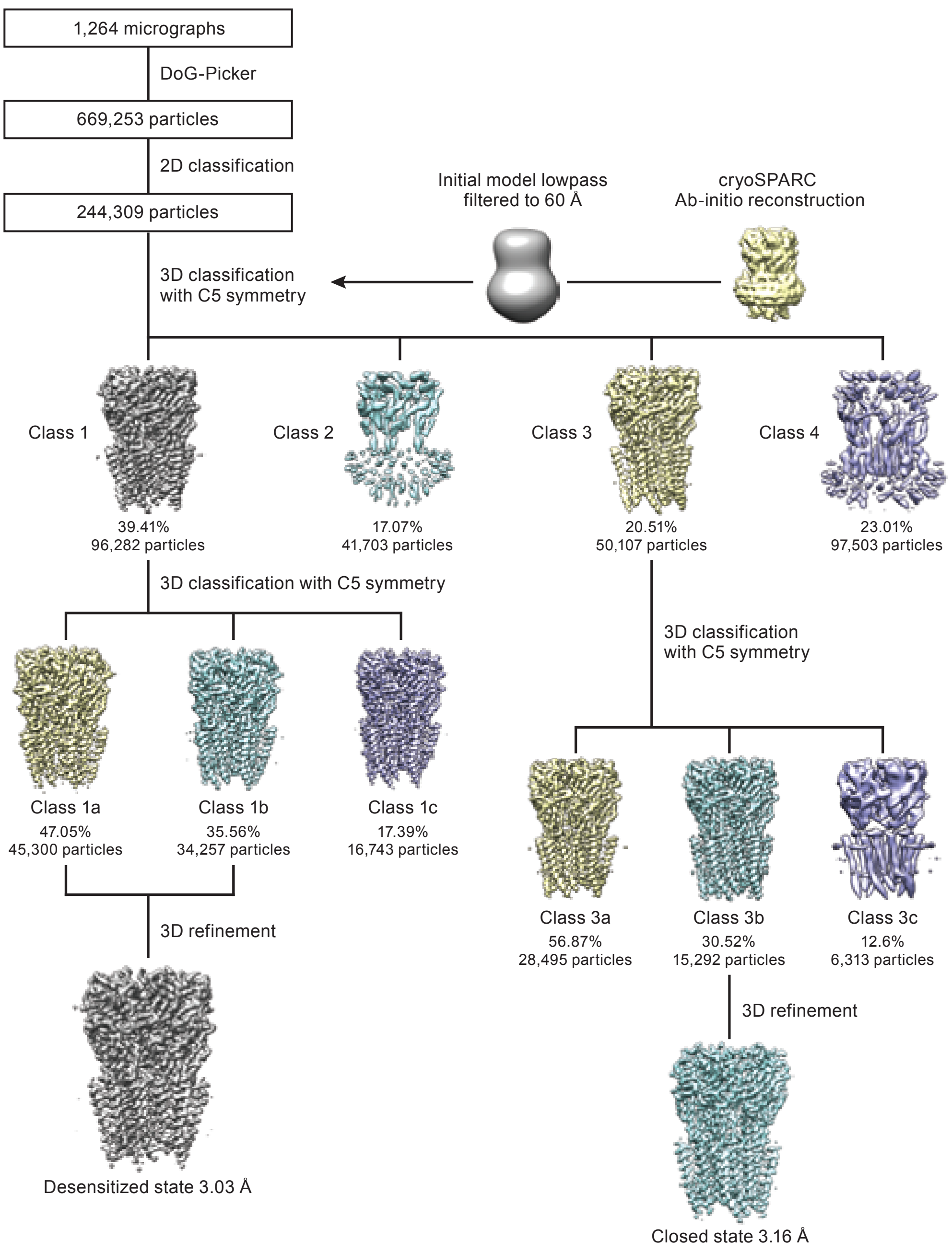


Supplementary Figure 1

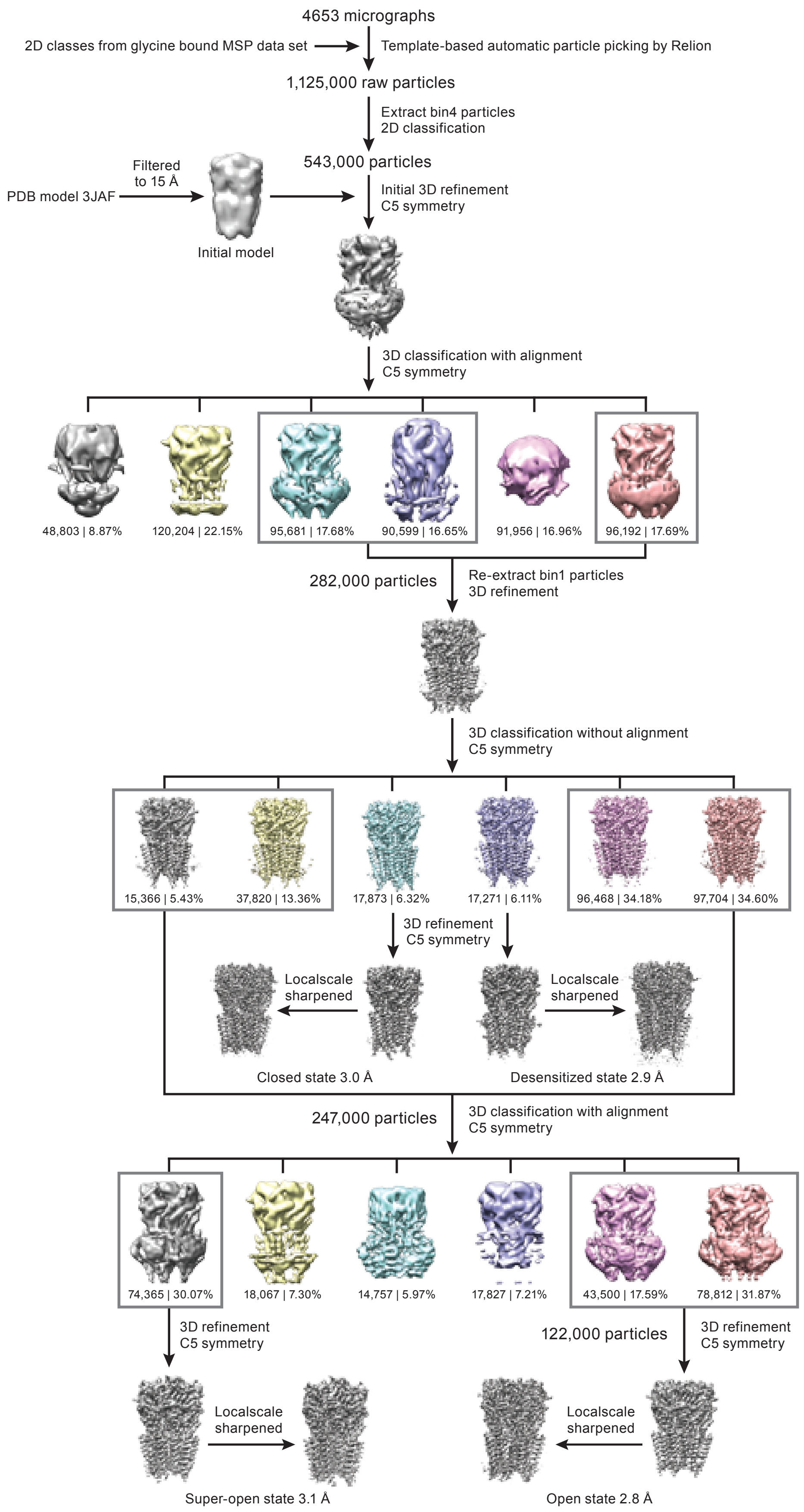


Supplementary Figure 2

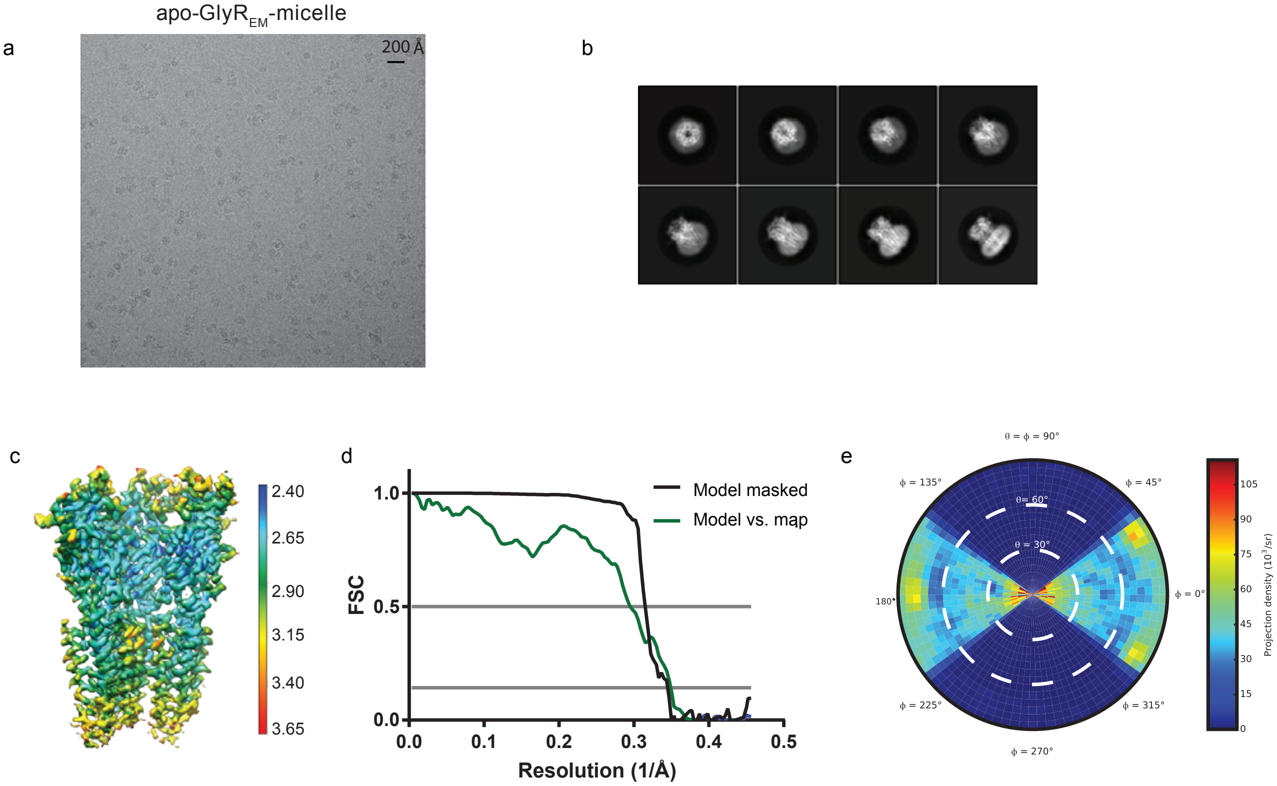


Supplementary Figure 3

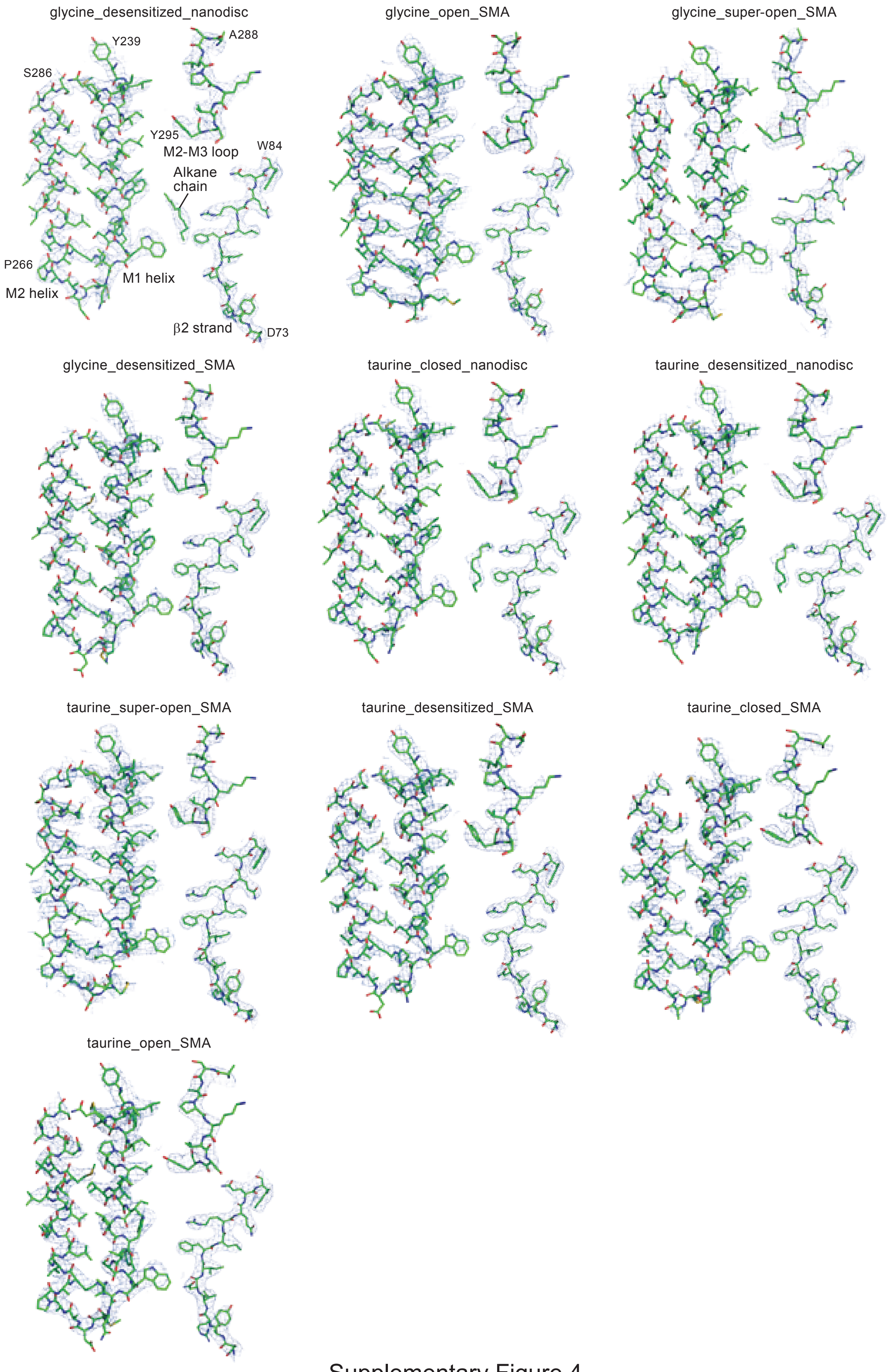


Supplementary Figure 4

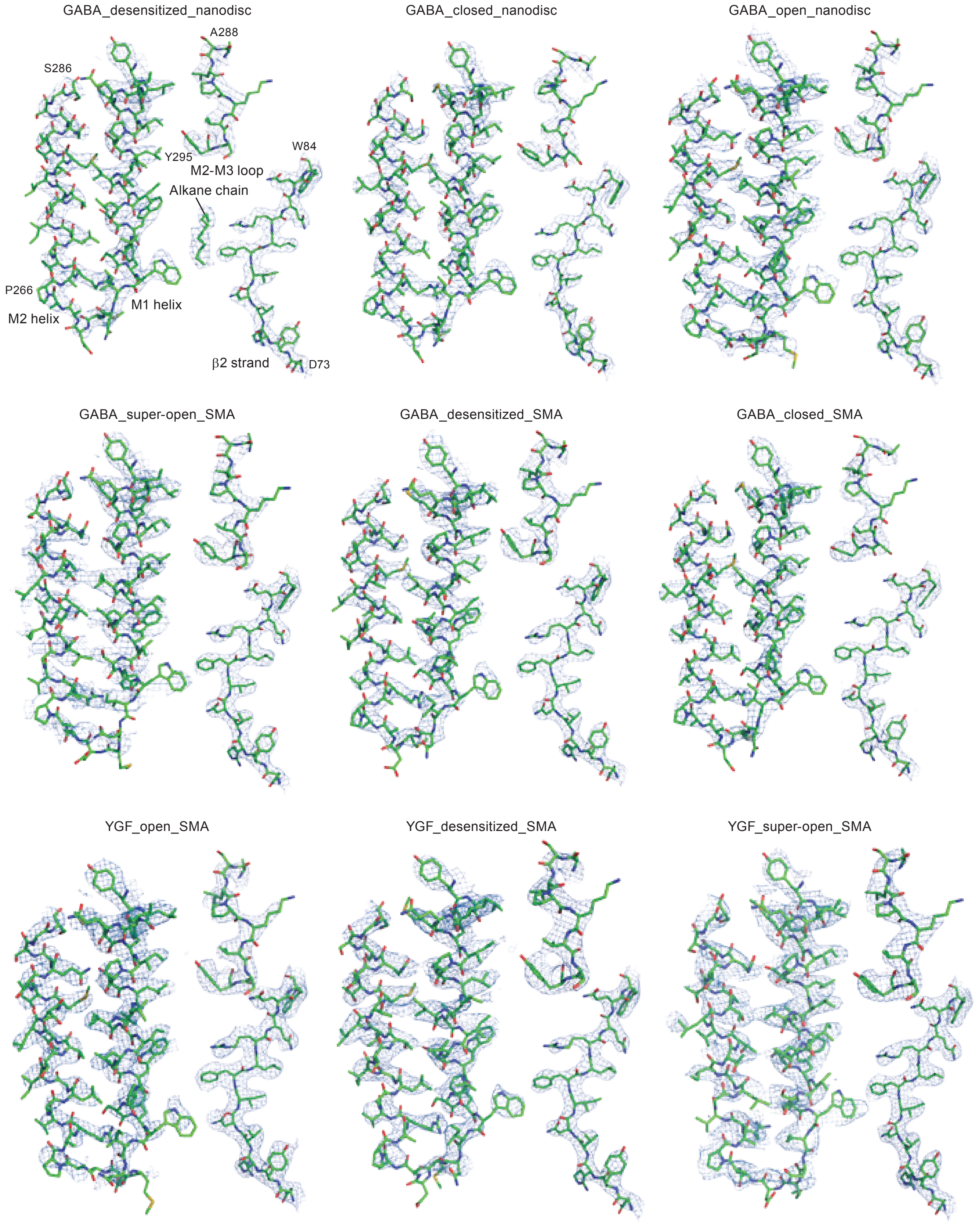


Supplementary Figure 5

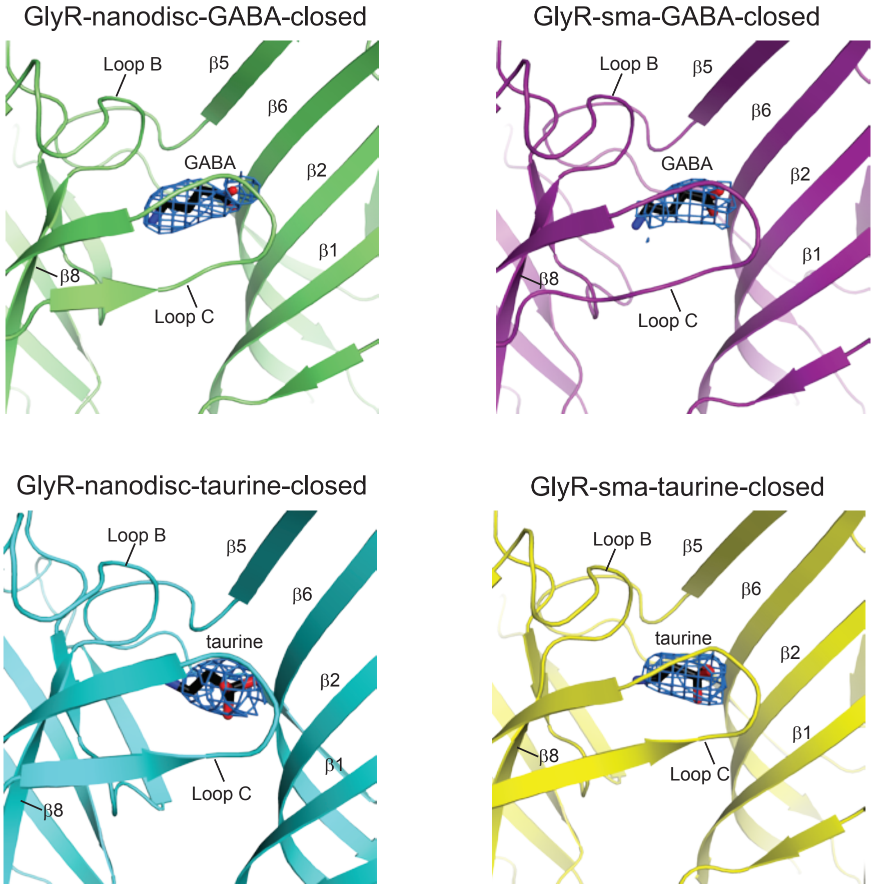


Supplementary Figure 6

**Supplementary Table 1 | Statistics for 3D reconstruction and model refinement in the nanodisc.**

| **Ligand**<br>**State** | **Glycine**<br>Desensitized<br>(EMD-20373)<br>(PDB 6PLR) | **Taurine**<br>Desensitized<br>(EMD-20374)<br>(PDB 6PLS) | **Taurine**<br>Closed<br>(EMD-20375)<br>(PDB 6PLT) | **GABA**<br>Desensitized<br>(EMD-20376)<br>(PDB 6PLU) | **GABA**<br>Closed<br>(EMD-20377)<br>(PDB 6PLV) |
|---|---|---|---|---|---|
| **Data collection and processing** | | | | | |
| Microscope | Arctica Talos | Arctica Talos | Arctica Talos | Arctica Talos | Arctica Talos |
| Camera | K2 Summit | K2 Summit | K2 Summit | K2 Summit | K2 Summit |
| Magnification | 45,000 | 45,000 | 45,000 | 45,000 | 45,000 |
| Voltage (kV) | 200 | 200 | 200 | 200 | 200 |
| Defocus range (μm) | -0.8 to -2.2 | -0.8 to -2.2 | -0.8 to -2.2 | -0.8 to -2.2 | -0.8 to -2.2 |
| Exposure time (s) | 10 | 10 | 10 | 10 | 10 |
| Dose rate ($e^-$/Å$^2$/s) | 6.1 | 6.1 | 6.1 | 6.1 | 6.1 |
| Number of frames (no.) | 100 | 100 | 100 | 100 | 100 |
| Pixel size (Å) | 0.899 | 0.899 | 0.899 | 0.899 | 0.899 |
| Symmetry imposed | C5 | C5 | C5 | C5 | C5 |
| Initial particles (no.) | 472910 | 669253 | 669253 | 493873 | 493873 |
| Final particles (no.) | 80121 | 124857 | 15292 | 39926 | 38383 |
| Map resolution (Å) | 3.2 | 3.0 | 3.2 | 3.3 | 3.3 |
| FSC threshold | 0.143 | 0.143 | 0.143 | 0.143 | 0.143 |
| Map resolution range (Å) | 3.0–3.4 | 2.7–3.9 | 3.1–3.9 | 3.0–3.8 | 2.9–4.1 |
| **Refinement** | | | | | |
| Initial model (PDB code) | 6PLS | 3JAF | 3JAD | 6PLS | 6PLT |
| Model resolution (Å) | 3.3 | 3.0 | 3.3 | 3.3 | 3.4 |
| FSC threshold | 0.5 | 0.5 | 0.5 | 0.5 | 0.5 |
| Model composition | | | | | |
| Non-hydrogen atoms | 14685 | 14610 | 14445 | 14665 | 14655 |
| Protein atoms | 13915 | 13805 | 13935 | 13835 | 13980 |
| Ligand atoms | 770 | 805 | 510 | 830 | 675 |
| *B* factors (Å$^2$) | | | | | |
| Protein | 83.0 | 48.9 | 120.0 | 84.3 | 69.1 |
| Ligand | 58.7 | 21.1 | 114.2 | 58.7 | 58.6 |
| Lipid | 88.1 | 67.5 | 96.4 | 94.8 | 66.3 |
| R.m.s. deviations | | | | | |
| Bond length (Å) | 0.004 | 0.008 | 0.005 | 0.008 | 0.009 |
| Bond angle (˚) | 0.837 | 1.027 | 0.835 | 0.952 | 0.996 |
| **Validation** | | | | | |
| Favored (%) | 95.96 | 97.41 | 96.72 | 97.06 | 97.48 |
| Allowed (%) | 4.04 | 2.59 | 3.28 | 2.94 | 2.52 |
| Disallowed (%) | 0 | 0 | 0 | 0 | 0 |
| Poor rotamers (%) | 0 | 0 | 0 | 0 | 0 |
| MolProbity score | 1.46 | 1.40 | 1.56 | 1.50 | 1.57 |
| Clash score | 4.26 | 5.21 | 4.00 | 5.55 | 4.11 |

**Supplementary Table 2 | Statistics for 3D reconstruction and model refinement of glycine and taurine bound GlyR in the SMA.**

| **Ligand**<br>**State** | **Glycine**<br>Desensitized<br>(EMD-20388)<br>(PDB 6PM5) | **Glycine**<br>Open<br>(EMD-20389)<br>(PDB 6PM6) | **Glycine**<br>Super-open<br>(EMD-20386)<br>(PDB 6PM4) | **Taurine**<br>Desensitized<br>(EMD-20383)<br>(PDB 6PM1) | **Taurine**<br>Open<br>(EMD-20384)<br>(PDB 6PM2) | **Taurine**<br>Super-open<br>(EMD-20382)<br>(PDB 6PM0) | **Taurine**<br>Closed<br>(EMD-20385)<br>(PDB 6PM3) |
|---|---|---|---|---|---|---|---|
| **Data collection and processing** | | | | | | | |
| Microscope | Titan Krios | Titan Krios | Titan Krios | Titan Krios | Titan Krios | Titan Krios | Titan Krios |
| Camera | K2 Summit | K2 Summit | K2 Summit | K2 Summit | K2 Summit | K2 Summit | K2 Summit |
| Magnification | 165,000 | 165,000 | 165,000 | 165,000 | 165,000 | 165,000 | 165,000 |
| Voltage (kV) | 300 | 300 | 300 | 300 | 300 | 300 | 300 |
| Defocus range (μm) | -1.0 to -2.0 | -1.0 to -2.0 | -1.0 to -2.0 | -1.0 to -2.0 | -1.0 to -2.0 | -1.0 to -2.0 | -1.0 to -2.0 |
| Exposure time (s) | 8 | 8 | 8 | 8 | 8 | 8 | 8 |
| Dose rate ($e^{-}$/Å$^2$/s) | 5.1 | 5.1 | 5.1 | 5.1 | 5.1 | 5.1 | 5.1 |
| Number of frames (no.) | 40 | 40 | 40 | 40 | 40 | 40 | 40 |
| Pixel size (Å) | 0.823 | 0.823 | 0.823 | 0.823 | 0.823 | 0.823 | 0.823 |
| Symmetry imposed | C5 | C5 | C5 | C5 | C5 | C5 | C5 |
| Initial particles (no.) | 1578324 | 1578324 | 1578324 | 1124737 | 1124737 | 1124737 | 1124737 |
| Final particles (no.) | 39586 | 155456 | 84978 | 17271 | 122322 | 74365 | 17873 |
| Map resolution (Å) | 3.1 | 2.9 | 4.0 | 3.0 | 2.8 | 3.1 | 3.0 |
| FSC threshold | 0.143 | 0.143 | 0.143 | 0.143 | 0.143 | 0.143 | 0.143 |
| Map resolution range (Å) | 2.5–4.4 | 2.5–4.5 | 3.4–5.5 | 2.5–4.0 | 2.5–4.5 | 2.7–4.3 | 2.7–3.9 |
| **Refinement** | | | | | | | |
| Initial model (PDB code) | 6PLS | 6PM5 | 6PM5 | 6PLS | 6PM5 | 6PM5 | 6PLT |
| Model resolution | 3.2 | 3.1 | 4.2 | 3.0 | 3.2 | 3.1 | 3.2 |
| FSC threshold | 0.5 | 0.5 | 0.5 | 0.5 | 0.5 | 0.5 | 0.5 |
| Model composition | | | | | | | |
| Non-hydrogen atoms | 14150 | 14100 | 14055 | 14160 | 14125 | 14080 | 14120 |
| Protein atoms | 13890 | 13840 | 13835 | 13930 | 13895 | 13850 | 13945 |
| Ligand atoms | 260 | 260 | 220 | 230 | 230 | 230 | 175 |
| *B* factors (Å$^2$) | | | | | | | |
| Protein | 97.8 | 101.4 | 242.1 | 94.1 | 98.7 | 111.5 | 48.8 |
| Ligand | 87.0 | 80.7 | - | 68.9 | 79.8 | 69.0 | 35.0 |
| R.m.s. deviations | | | | | | | |
| Bond length (Å) | 0.001 | 0.005 | 0.007 | 0.007 | 0.008 | 0.007 | 0.005 |
| Bond angle (˚) | 0.399 | 0.911 | 0.927 | 0.954 | 1.021 | 0.968 | 0.916 |
| **Validation** | | | | | | | |
| Favored (%) | 97.66 | 95.96 | 90.12 | 96.49 | 95.61 | 95.91 | 96.95 |
| Allowed (%) | 2.34 | 4.04 | 9.88 | 3.51 | 4.39 | 4.09 | 3.05 |
| Disallowed (%) | 0 | 0 | 0 | 0 | 0 | 0 | 0 |
| Poor rotamers | 0 | 0 | 0 | 0 | 0 | 0 | 0 |
| MolProbity score | 1.35 | 1.61 | 1.97 | 1.63 | 1.52 | 1.55 | 1.58 |
| Clash score | 5.03 | 5.13 | 7.69 | 5.69 | 4.47 | 5.03 | 4.67 |

**Supplementary Table 3 | Statistics for 3D reconstruction and model refinement of the GABA bound GlyR in the SMA.**

| **Ligand**<br>**State** | **GABA**<br>Desensitized<br>(EMD-20379)<br>(PDB 6PLX) | **GABA**<br>Open<br>(EMD-20380)<br>(PDB 6PLY) | **GABA**<br>Super-open<br>(EMD-20378)<br>(PDB 6PLW) | **GABA**<br>Closed<br>(EMD-20381)<br>(PDB 6PLZ) |
|---|---|---|---|---|
| **Data collection and processing** | | | | |
| Microscope | Titan Krios | Titan Krios | Titan Krios | Titan Krios |
| Camera | K2 Summit | K2 Summit | K2 Summit | K2 Summit |
| Magnification | 165,000 | 165,000 | 165,000 | 165,000 |
| Voltage (kV) | 300 | 300 | 300 | 300 |
| Defocus range (μm) | -1.0 to -2.0 | -1.0 to -2.0 | -1.0 to -2.0 | -1.0 to -2.0 |
| Exposure time (s) | 8 | 8 | 8 | 8 |
| Dose rate ($e^{-}$/Å$^2$/s) | 5.1 | 5.1 | 5.1 | 5.1 |
| Number of frames | 40 | 40 | 40 | 40 |
| Pixel size (Å) | 0.823 | 0.823 | 0.823 | 0.823 |
| Symmetry imposed | C5 | C5 | C5 | C5 |
| Initial particles (no.) | 2062519 | 2062519 | 2062519 | 2062519 |
| Final particles (no.) | 20845 | 150199 | 121470 | 31667 |
| Map resolution (Å) | 2.9 | 2.9 | 3.0 | 3.0 |
| FSC threshold | 0.143 | 0.143 | 0.143 | 0.143 |
| Map resolution range (Å) | 2.6–4.2 | 2.7–4.3 | 2.7–4.3 | 2.6–4.2 |
| **Refinement** | | | | |
| Initial model (PDB code) | 6PLS | 6PM5 | 6PM5 | 6PLT |
| Model resolution (Å) | 3.1 | 3.1 | 3.1 | 3.3 |
| FSC threshold | 0.5 | 0.5 | 0.5 | 0.5 |
| Model composition | | | | |
| Non-hydrogen atoms | 14240 | 14165 | 14095 | 14210 |
| Protein atoms | 13970 | 13935 | 13865 | 13980 |
| Ligand atoms | 270 | 230 | 230 | 230 |
| *B* factors (Å$^2$) | | | | |
| Protein | 110.6 | 127.2 | 139.3 | 92.0 |
| Ligand | 151.0 | 107.5 | 104.6 | 90.1 |
| R.m.s. deviations | | | | |
| Bond length (Å) | 0.006 | 0.005 | 0.010 | 0.006 |
| Bond angle (˚) | 0.953 | 0.865 | 0.965 | 0.929 |
| **Validation** | | | | |
| Favored (%) | 96.20 | 94.91 | 96.49 | 96.01 |
| Allowed (%) | 3.80 | 5.09 | 3.51 | 3.99 |
| Disallowed (%) | 0 | 0 | 0 | 0 |
| Poor rotamers | 0 | 0 | 0 | 0 |
| MolProbity score | 1.53 | 1.62 | 1.50 | 1.59 |
| Clash score | 6.04 | 5.12 | 5.06 | 4.54 |

**Supplementary Table 4 | Statistics for 3D reconstruction and model refinement of YGF mutant and apo-GlyR$_{EM}$ in the micelle.**

| **Ligand**<br>**State** | **GABA**<br>Open<br>(EMD-20370)<br>(PDB 6PLO) | **GABA**<br>Super-open<br>(EMD-20372)<br>(PDB 6PLQ) | **GABA**<br>Desensitized<br>(EMD-20371)<br>(PDB 6PLP) | **-**<br>Apo-GlyR$_{EM}$<br>(EMD-20518)<br>(PDB 6PXD) |
|---|---|---|---|---|
| **Data collection and processing** | | | | |
| Microscope | Titan Krios | Titan Krios | Titan Krios | Titan Krios |
| Camera | K3 BioQuantum | K3 BioQuantum | K3 BioQuantum | K2 Summit |
| Magnification | 130,000 | 130,000 | 130,000 | 130,000 |
| Voltage (kV) | 300 | 300 | 300 | 300 |
| Defocus range (μm) | -1.0 to -2.0 | -1.0 to -2.0 | -1.0 to -2.0 | -1.2 to -2.5 |
| Exposure time (s) | 1.5 | 1.5 | 1.5 | 6 |
| Dose rate ($e^{-}$/Å$^2$/s) | 18.8 | 18.8 | 18.8 | 9.2 |
| Number of frames | 60 | 60 | 60 | 30 |
| Pixel size (Å) | 0.648 | 0.648 | 0.648 | 1.040 |
| Symmetry imposed | C5 | C5 | C5 | C5 |
| Initial particles (no.) | 1808565 | 1808565 | 1808565 | 204323 |
| Final particles (no.) | 32386 | 42097 | 27496 | 115864 |
| Map resolution (Å) | 3.3 | 3.4 | 3.3 | 2.9 |
| FSC threshold | 0.143 | 0.143 | 0.143 | 0.143 |
| Map resolution range (Å) | 3.2–4.4 | 3.0–4.1 | 3.1–4.3 | 2.4–3.7 |
| **Refinement** | | | | |
| Initial model (PDB code) | 6PLY | 6PLW | 6PLX | 3JAD |
| Model resolution (Å) | 3.4 | 3.5 | 3.4 | 3.2 |
| FSC threshold | 0.5 | 0.5 | 0.5 | 0.5 |
| Model composition | | | | |
| Non-hydrogen atoms | 14165 | 14110 | 14240 | 13760 |
| Protein atoms | 13935 | 13880 | 14010 | 1616 |
| Ligand atoms | 230 | 230 | 230 | 144 |
| *B* factors (Å$^2$) | | | | |
| Protein | 131.6 | 150.6 | 109.8 | 69.4 |
| Ligand | 116.7 | 131.7 | 94.0 | - |
| R.m.s. deviations | | | | |
| Bond length (Å) | 0.007 | 0.007 | 0.001 | 0.007 |
| Bond angle (˚) | 0.999 | 0.977 | 0.416 | 1.30 |
| **Validation** | | | | |
| Favored (%) | 95.44 | 94.91 | 97.95 | 95.27 |
| Allowed (%) | 4.56 | 5.09 | 2.05 | 4.73 |
| Disallowed (%) | 0 | 0 | 0 | 0 |
| Poor rotamers | 0 | 0 | 0 | 1.32 |
| MolProbity score | 1.34 | 1.46 | 1.65 | 1.99 |
| Clash score | 6.08 | 3.13 | 6.08 | 11.24 |

**Supplementary Table5 | Resolution summary for all the maps.**

| Membrane mimic | Agonist | State | Resolution (Å) |
|---|---|---|---|
| MSP-lipid nanodisc | glycine | Desensitized | 3.2 |
| | taurine | Desensitized | 3.0 |
| | | Closed | 3.2 |
| | GABA | Desensitized | 3.3 |
| | | Closed | 3.3 |
| SMA | glycine | Open | 2.9 |
| | | Desensitized | 3.1 |
| | | Super-open | 4.0 |
| | taurine | Closed | 3.0 |
| | | Open | 2.8 |
| | | Desensitized | 3.0 |
| | | Super-open | 3.1 |
| | GABA | Closed | 3.0 |
| | | Open | 2.9 |
| | | Desensitized | 2.9 |
| | | Super-open | 3.0 |
| | YGF-GABA | Open | 3.3 |
| | | Super-open | 3.4 |
| | | Desensitized | 3.3 |
| Micelle | - | Apo-$GlyR_{EM}$ | 2.9 |

**Supplementary Table 6 | Binding pocket metrics for Cys-loop family.†**

| | $d_x$ (Å) | $V_{ligand}$ (Å$^3$) | $D_{F175\text{-}S145}$(Å) | $D_{T220\text{-}R81}$(Å) | |
|---|---|---|---|---|---|
| **Glycine receptor** | | | | | |
| Glycine(6PM6) | 2.7 | 53.1 | 9.1 | 9.3 | Full Agonist |
| Taurine(6PM2) | 3.5 | 79.0 | 9.5 | 9.4 | Partial Agonist |
| GABA(6PLY) | 4.1 | 86.5 | 9.6 | 9.6 | Partial Agonist |
| Strychnine(3JAD) | 3.1 | 258.4 | 10.4 | 12.1 | Antagonist |
| Apo(5WP7) | | | 10.0 | 12.6 | |
| Efficacy: glycine > taurine > GABA > strychnine | | | | | |
| **Nicotinic acetylcholine receptor** | | | | | |
| Acetylcholine(3WIP)[1] | 1.9* | 120 | 9.2 | 8.8 | Full Agonist |
| Nicotine(5KXI)[2] | 2.2* | 137 | 9.4 | 9.8 | Partial Agonist |
| Varenicline(4AFT)[3] | 2.3 | 167 | 9.0 | 11.2 | Partial Agonist |
| Efficacy: Acetylcholine>Nicotine >Varenicline | | | | | |
| **5HT3 receptor** | | | | | |
| 5HT3(6HIO)[4] | 2.3 | 138.6 | 7.4 | 13.4 | Full Agonist |
| Varenicline(5AIN)[5] | 2.6 | 167.3 | 9.5 | 16.4 | Partial Agonist |
| Tropisetron(6HIS)[4] | 2.1 | 221.0 | 8.1 | 15.6 | Antagonist |
| Apo (6BE1)[6] | | | 8.6 | 15.7 | |
| Efficacy: Serotonin >Varenicline> Tropisetron | | | | | |
| **GABAA receptor** | | | | | |
| GABA(6DW0)[7] | 3.7 | 86.5 | 9.3 | 12.0 | Full Agonist |
| BCC(6HUK)[8] | 3.2 | 263.5 | 10.1 | 12.2 | Antagonist |
| Efficacy: GABA > BCC | | | | | |

* values are obtained by Tripathy et al.[9].

† Four measurements are performed for each structure: 1) the $d_x$ (in Å) as defined by Auerbach and colleagues[9]; 2) the volume of ligand ($V_{ligand}$; in Å$^3$) calculated using the Molecular Volume Calculator[10]; 3) the distance between the Cα atoms of T175 (+) and S145 (-) ($D_{F175\text{-}S145}$; in Å) in GlyR or the corresponding Cα atoms in the non-GlyR structures; 4) the distance between the Cα atoms from T220 (+) and R81 (-) ($D_{T220\text{-}R81}$; in Å) in GlyR or the corresponding Cα atoms in the non-GlyR structures. PDB codes are noted in parentheses.
